# Changes in flexibility but not in compactness underlie the thermal adaptation of prokaryotic adenylate kinases

**DOI:** 10.1101/2024.09.04.611173

**Authors:** Dimitrios - Georgios Kontopoulos, Ilias Patmanidis, Timothy G. Barraclough, Samraat Pawar

## Abstract

Understanding the structural changes that enable enzymes to remain active in extreme thermal conditions is of broad scientific interest for both fundamental and applied biological research. Three key mechanisms that underlie the thermal adaptation of enzymes are modifications in structural flexibility, compactness, and the contacts formed among amino acids. However, most previous studies on these topics have been limited to small sample sizes or a narrow taxonomic focus, and the importance of these factors to thermal adaptation remains poorly understood. In this study, we combined molecular dynamics simulations and phylogenetic comparative analyses to thoroughly analyse the structural factors underlying thermal adaptation in adenylate kinase—a key enzyme involved in cellular energy balance and homeostasis—across 70 prokaryotic species. We detect systematic increases in the flexibility of the enzyme with temperature, both across and within species. In contrast, structural compactness appears to be almost completely independent of temperature. Finally, we uncover a remarkable diversity in the number and types of amino acid contacts observed in different adenylate kinases that cannot be explained solely by temperature. Our results suggest that there are multiple paths toward the adaptation of prokaryotic adenylate kinases to extreme thermal environments and that these paths are generally accessible through changes in flexibility.

**Lay summary:** The structure of a given enzyme can vary considerably among species, reflecting local environmental conditions to an extent. To this day, we do not have a clear picture of the impacts of the thermal environment on enzyme structure. To fill this gap, we performed a structural comparison of the enzyme adenylate kinase (ADK) from 70 species of bacteria and archaea. We find that rises in temperature tend to increase the flexibility of the enzyme. However, at any given temperature, ADKs from cold environments tend to be more flexible than those from hot environments. In contrast, the compactness of the enzyme did not vary consistently with temperature. Finally, we found that the pattern of amino acid contacts can vary dramatically across ADKs of different species, in a manner that cannot be predicted by temperature alone. Overall, our study shows that there are multiple ways to evolve an enzyme structure that can tolerate extreme temperatures, with a key constraint being maintaining sufficient flexibility at temperatures typically experienced by each species.

## Introduction

Changes in environmental temperature directly or indirectly affect most biological processes, from the activity and stability of enzymes to the functioning of entire ecosystems (Pawar et al., 2015; Clarke, 2017; Vázquez et al., 2017; Knapp and Huang, 2022). Understanding the mechanisms through which biological systems respond to temperature allows us not only to predict their future dynamics in the face of climate change but also to identify ways to modify them for ecological, environmental, or biotechnological purposes.

At the level of individual enzymes, both optimal temperature (where activity is maximised) and melting temperature (where half of the enzyme population is in the unfolded state) have been shown to correlate strongly and positively with growth temperature in microbes (Engqvist, 2018; Stark et al., 2022).

This implies that the thermal adaptation of microbial growth rate should depend on concordant changes in the stability and activity of key enzymes, achieved through modifications of their three-dimensional structures (Feller, 2010).

The most commonly discussed hypothesis for the adaptation of enzyme structures to different thermal environments is the “corresponding states” hypothesis, introduced by Somero (1978). This hypothesis posits that orthologous enzymes from different thermal environments should have similar kinetic and thermodynamic properties at their respective native temperatures. One such property is global structural flexibility which, for a given enzyme, should increase linearly with temperature due to the increase in kinetic energy. Because of this, the corresponding states hypothesis necessarily implies that, at a common (normalisation) temperature, cold-adapted enzyme orthologs should be more flexible than warm-adapted orthologs (Fig. 1).

**Fig. 1:**
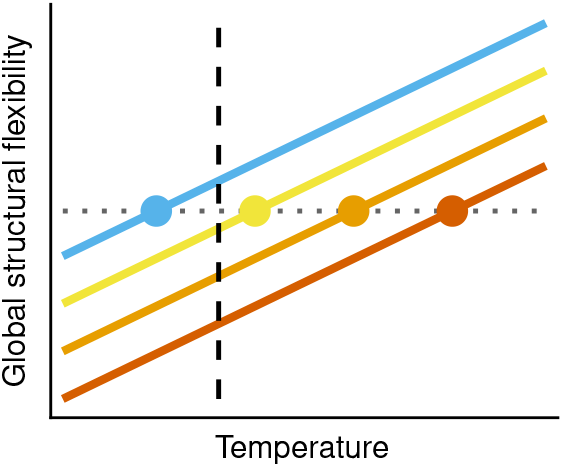
The relationship between the global structural flexibility of a given enzyme and temperature, according to the corresponding states hypothesis. The four colours represent orthologs of the same enzyme from different thermal environments. The corresponding states hypothesis posits that if flexibility is compared at the native temperature of each enzyme (circles), there should be no variation among orthologs. In other words, the interspecific slope (horizontal dotted line) should be zero. In contrast, if the four orthologs are compared at a common temperature (dashed vertical line), the coldest-adapted ortholog (blue) should be the most flexible.

A second, complementary hypothesis for structural changes that may underlie thermal adaptation of enzymes is an increase in structure compactness at high-temperature environments (Zhu et al., 1998; Robinson-Rechavi et al., 2006; Tompa et al., 2016). As enzyme structure becomes more compact, especially at its surface, its exposure to high-temperature solvents is reduced, decreasing the probability of denaturation. High structural compactness may also facilitate the formation of contacts among amino acid side chains—such as salt bridges—which can further stabilize the structure, making the denatured state less energetically favourable (Tompa et al., 2016).

Previous studies have found mixed support for the aforementioned hypotheses (e.g., Karshikoff and Ladenstein 1998; Szilágyi and Závodszky 2000; Fitter et al. 2001; Collins et al. 2003; Hajdú et al. 2008; Cipolla et al. 2012; Nguyen et al. 2017; Maffucci et al. 2020; Diessner et al. 2024; Jenney Jr et al. 2024), but such studies generally suffer from certain common limitations. In particular, because of experimental and/or computational constraints, most studies tend to compare only a small number of orthologs—often as low as two—which are usually separated by long phylogenetic distances. This makes it difficult to disentangle differences among orthologs that arose through thermal adaptation from those that arose randomly over many millions of years of evolution (see Uyeda et al. 2018). Due to small sample sizes, few studies report quantitative estimates (i.e., intercepts and slopes) of the impacts of temperature on key structural variables across and/or within enzymes from different species. Such estimates could be directly integrated into data syntheses, allowing for systematic comparisons of the effects of temperature across different types of enzymes. On the other hand, studies with larger sample sizes often suffer from the opposite problem, that is, they tend to focus on relatively narrow taxonomic groups and, therefore, it is often not possible to evaluate the generality of the conclusions drawn.

To thoroughly examine the structural changes involved in thermal adaptation of enzymes across a sufficiently wide taxonomic range, here we study the essential enzyme adenylate kinase (ADK). ADK catalyses the reversible conversion of ATP and AMP to ADP. Balancing the levels of these three nucleotides is crucial for cellular homeostasis and metabolic versatility, and therefore, the ADK activity strongly determines organismal fitness (Couñago and Shamoo, 2005; Couñago et al., 2006, 2008; Peña et al., 2010; Tükenmez et al., 2016).

ADKs have three major structural domains: i) the CORE domain which comprises the bulk of the structure, ii) the NMPbind domain where AMP is bound, and iii) the LID domain where ATP is bound (Li et al. 2015; Fig. 2). The CORE domain is rigid, whereas the two other domains are flexible, allowing the structure to shift between catalytically-inactive (open; e.g., Fig. 2A) and active (closed; Fig. 2B) conformations (Kovermann et al., 2017). Across prokaryotes, there are two main sources of variation in the ADK structure. First, instead of a long LID domain (Fig. 2A,B), some prokaryotic ADKs (e.g., that of *Mycobacterium tuberculosis*; Fig. 2C) have a short LID (Bellinzoni et al., 2006). Second, while most prokaryotic ADKs are monomeric, some archaeal ADKs (e.g., that of *Sulfolobus acidocaldarius*; Fig. 2D) have an additional beta hairpin which enables them to operate in homotrimeric rather than monomeric form (Vonrhein et al., 1998).

**Fig. 2:**
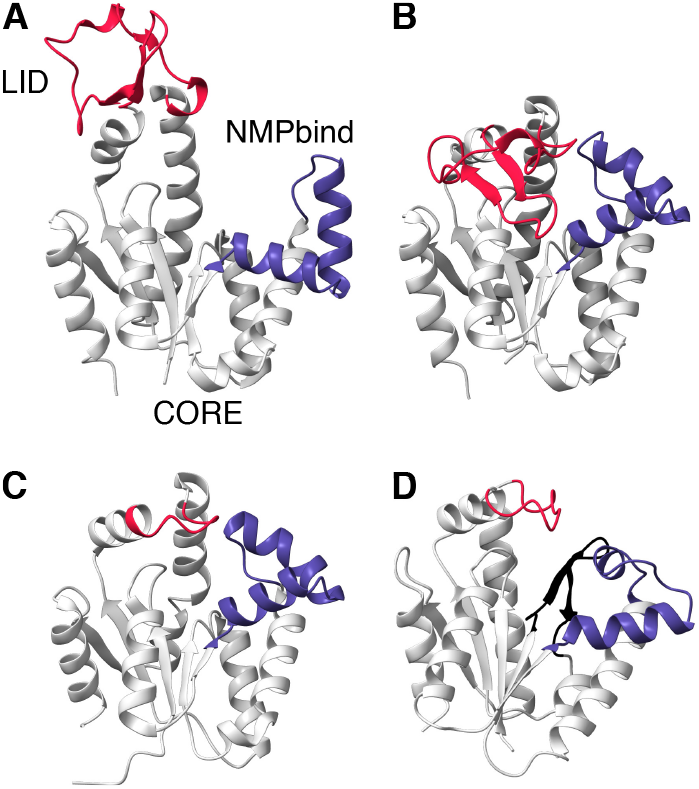
Main structural characteristics of prokaryotic ADKs. (A-B) Open and closed conformations of the ADK of the bacterium *Streptococcus pneumoniae* (PDB accession codes: 4NTZ and 4NU0). This ADK is monomeric and has a long LID domain (in red), in contrast to the monomeric ADK of the bacterium *Mycobacterium tuberculosis* (C; PDB accession code: 2CDN), whose LID domain is short. (D) The monomer of the trimeric ADK of the archaeon *Sulfolobus acidocaldarius* (PDB accession code: 1NKS). This ADK has a short LID domain and an additional beta hairpin (in black). The figure was generated with UCSF ChimeraX (v. 1.7.1; Pettersen et al. 2021).

Using ADK as our study system, here we ask three main questions:

1. Do ADKs from different thermal environments occupy distinct regions of the structural parameter space?
2. Does environmental temperature systematically influence structural flexibility and compactness at the intra- and/or interspecific levels?
3. Does adaptation of ADKs to extreme thermal environments require the formation (or absence) of several specific amino acid contacts?

To address these questions, we manually curated a dataset of 70 ADK structures from diverse bacterial and archaeal species, and thermal environments. We first performed a phylogenetic analysis to evaluate the extent of amino acid sequence conservation among ADK orthologs. Next, we conducted molecular dynamics simulations to obtain estimates of flexibility, compactness, and presence of amino acid contacts at various temperatures. Finally, we quantified the relationships between these factors and temperature through phylogenetic comparative methods.

## Results and Discussion

### Dataset of prokaryotic ADKs and phylogenetic distribution

We classified ADKs in our dataset based on the thermal preferences of their corresponding species as a) psychrophilic (optimal growth temperature up to 20^*°*^C), b) mesophilic (optimal growth temperature between 20 and 40^*°*^C), c) thermophilic (optimal growth temperature between 40 and 80^*°*^C), or d) hyperthermophilic (optimal growth temperature higher than 80^*°*^C). Our dataset includes ADKs from 8 psychrophiles, 38 mesophiles, 16 thermophiles, and 8 hyperthermophiles. The LID domain is long in 49 and short in 21 ADKs, whereas 12 out of 70 ADKs have the beta hairpin that allows them to form homotrimers.

Mapping these thermal groups on the species’ phylogeny (see Methods) revealed a scattered phylogenetic distribution (Fig. 3A), which should provide adequate statistical power for addressing our three main questions. In particular, the phylogeny includes 8 pairs of sister species (species immediately descending from a common node) that belong to different thermal groups. Neither the ADK type (monomeric or trimeric) nor the length of the LID are monophyletically distributed, possibly due to interspecific recombination events, similar to those previously reported for ADKs from the bacterial genus *Neisseria* (Feil et al., 1995, 1996).

**Fig. 3:**
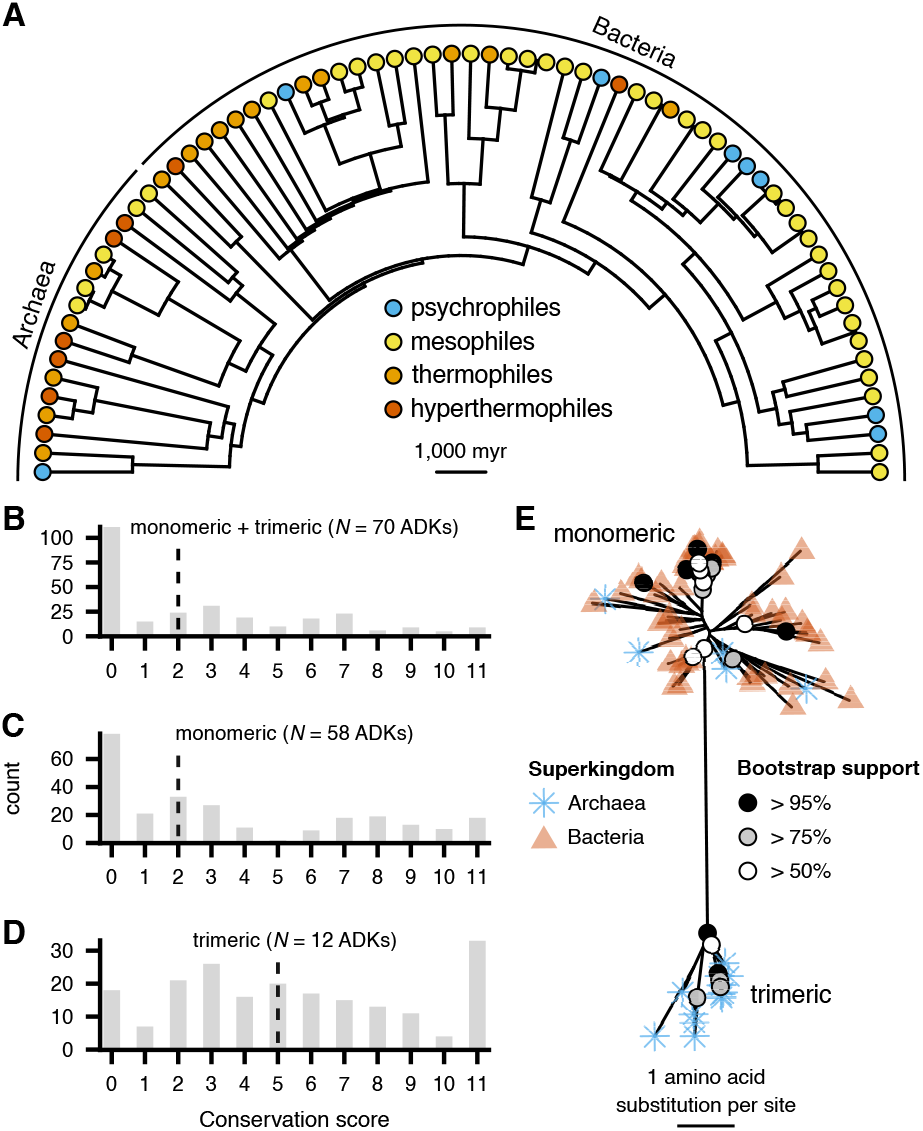
Overview of the 70 prokaryotic ADKs included in this study. (A) Distribution of the four thermal groups across the species’ phylogeny, visualized using the phytools R package (v.2.1-1; Revell 2024). (B-D) Conservation scores for each column of the alignments of ADK amino acid sequences, obtained with Jalview (v. 2.11.3.3; Waterhouse et al. 2009). A value of 11 indicates columns where all sequences share a common amino acid, whereas lower values correspond to lower degrees of physicochemical conservation (Livingstone and Barton, 1993). The dashed vertical line in each panel stands for the median value. (E) The maximum-likelihood gene tree of all 70 ADK sequences, as estimated with IQ-TREE (v. 2.3.5; Minh et al. 2020) and visualized with the ggtree R package (v. 3.12.0; Yu et al. 2017). Branches with bootstrap support values greater than 50% are explicitly annotated.

### Amino acid sequence conservation

We next evaluated the degree of amino acid sequence conservation in each column of the alignment of a) all ADKs, b) monomeric ADKs, and c) trimeric ADKs (Fig. 3B-D). We found evidence for low conservation among most alignment columns, but especially for monomeric ADKs (Fig. 3C). The sequences of trimeric ADKs appeared to be relatively more conserved (Fig. 3D) which may be due to the need to maintain the trimerization interface. A gene tree reconstructed from the alignment of all 70 ADK amino acid sequences revealed two major clusters, one for monomeric and one for trimeric ADKs (Fig. 3E). Most branches of the gene tree had very low bootstrap support, which is in line with the fact that most alignment columns were weakly conserved.

### Comparison of the structural parameter space occupied by the four thermal groups

To understand whether thermal adaptation of prokaryotic ADKs involves dramatic shifts in the structural parameter space (our first main question), we conducted 10 replicate molecular dynamics simulations for a combined length of 2 *µ*s per ADK (see Methods). In these simulations, we specified a different (“native”) temperature for each group, namely 280 K (6.85^*°*^C) for psychrophilic ADKs, 300 K (26.85^*°*^C) for mesophilic ADKs, 330 K (56.85^*°*^C) for thermophilic ADKs, or 355 K (81.85^*°*^C) for hyperthermophilic ADKs. The simulations yielded estimates of 7 structural variables that describe the flexibility, compactness, and presence/lack of amino acid contacts per ADK (see the caption of Fig. 4 for a description of each variable), which we analysed through phylogenetic principal components analysis (pPCA).

**Fig. 4:**
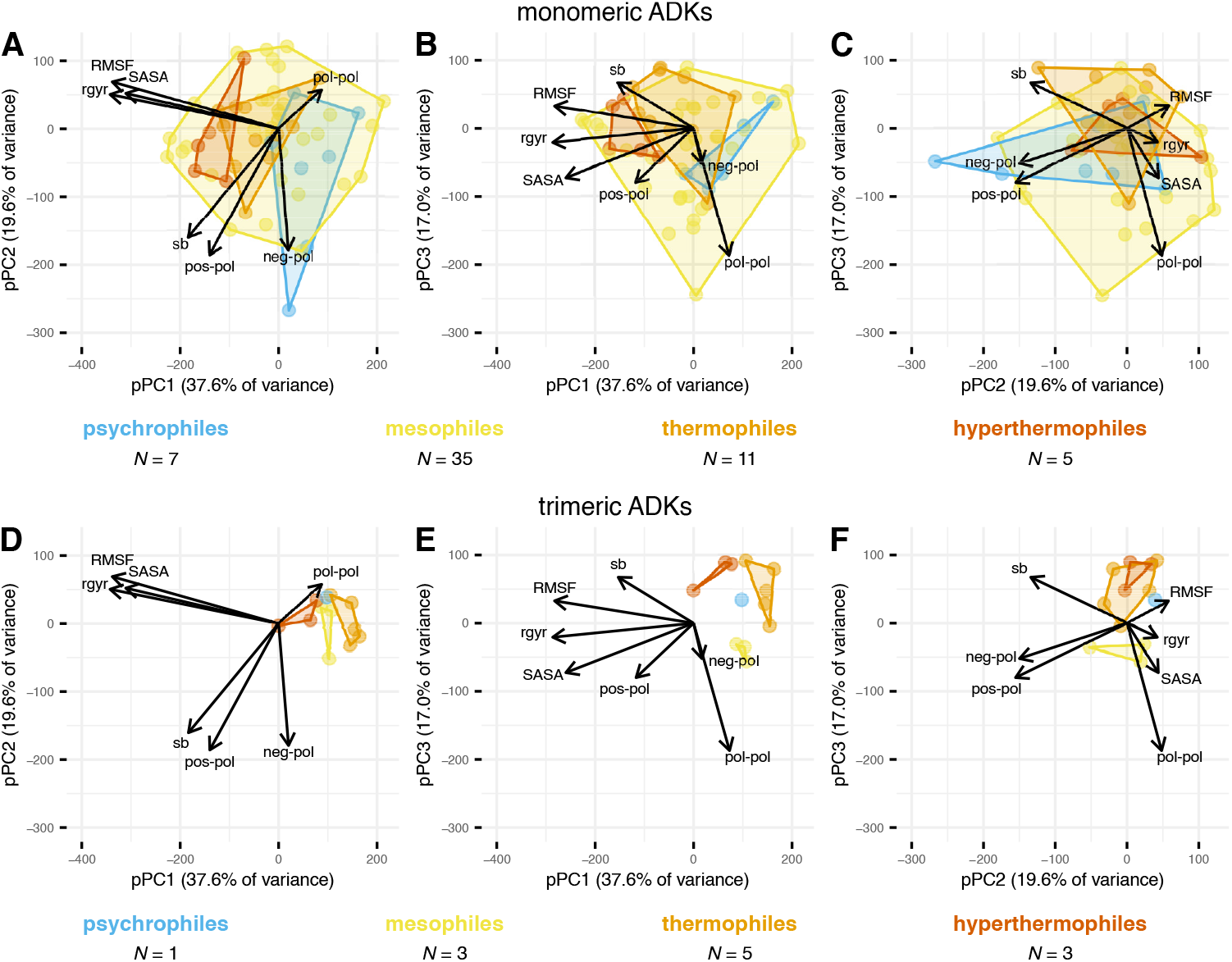
Phylogenetic principal components analysis of 7 structural variables (arrows) for the 70 prokaryotic ADKs of the present study. RMSF stands for the root mean square fluctuation of alpha carbon atoms and is a measure of structural flexibility. SASA is the solvent-accessible surface area, a proxy of surface compactness. r_gyr_ is the radius of gyration of alpha carbon atoms and is a measure of overall structure compactness. The remaining variables represent counts of different contact types between pairs of amino acid side chains: i) negatively charged - polar (“neg-pol”), ii) positively charged - polar (“pos-pol”), iii) polar - polar (“pol-pol”), or iv) salt bridges (“sb”; contacts between a negatively charged and a positively charged side chain). The four thermal groups are shown in different colours, whereas monomeric (A-C) and trimeric ADKs (D-F) are shown separately for clarity.

Overall, the four thermal groups overlapped considerably throughout the parameter space, with the most pronounced separation observed in the first phylogenetic principal component (pPC1) where psychrophiles and hyperthermophiles had no overlap (Fig. 4). Nevertheless, mesophiles occurred throughout the entire range of pPC1, with values exceeding those of both psychrophiles and hyperthermophiles. The variables that most strongly correlated with pPC1 reflected structural flexibility (RMSF) and compactness (SASA and r_gyr_). Monomeric ADKs (Fig. 4A) covered a much wider range of parameter space than their trimeric orthologs (Fig. 4B), which may again reflect the structural constraints that trimerization imposes. These results suggest that a) there are likely multiple evolutionary paths towards adaptation to extreme thermal environments, b) such environments may be well tolerated by certain mesophilic ADKs, and that c) changes in flexibility and/or compactness may indeed contribute to the thermal adaptation of prokaryotic ADKs.

### Impacts of temperature on ADK flexibility

The simulations performed so far would enable us to infer the interspecific relationship between ADK flexibility (the RMSF variable) and temperature. To also capture the intraspecific relationship for our second main question, we performed additional simulations for five ADKs at seven more temperatures ranging from 6.85^*°*^C to 94.35^*°*^C (henceforth referred to as “non-native temperatures”). Given that monomeric ADKs occupy a much wider range of parameter space (Fig. 4), the five ADKs that we selected were all monomeric. We then fitted a series of candidate models to simultaneously estimate the intra- and interspecific relationships (if any) between RMSF and temperature. In these models, we accounted for a) the uncertainty around each RMSF estimate, b) the ADK type (monomeric or trimeric), c) the LID length (short or long), and/or d) the evolutionary relationships among species (through a phylogenetic random effect on the intercept). The best-supporting model was identified through model selection (see Methods).

At the within-species level, we found that flexibility (RMSF values) consistently increased with temperature (Fig. 5A and Supplementary Fig. S4), as expected from the increase in the total energy of the system at higher temperatures. On top of this, the interspecific temperature slope was not zero (as the corresponding states hypothesis predicts; Fig. 1) but positive, with long-LID and short-LID ADKs differing in their intercepts. The aforementioned factors explained 64% of variance in RMSF 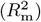, whereas a phylogenetic random effect on the intercept captured an additional 20% of variance 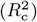. This suggests that evolutionary shifts in flexibility can also arise in response to other—phylogenetically-structured—factors (e.g., the range of pH values tolerated by each species). In particular, while almost all archaea in our dataset appear to have evolved low temperature- and LID-corrected RMSF values, bacterial clades are split between high- and low-RMSF groups (Fig. 5B).

**Fig. 5:**
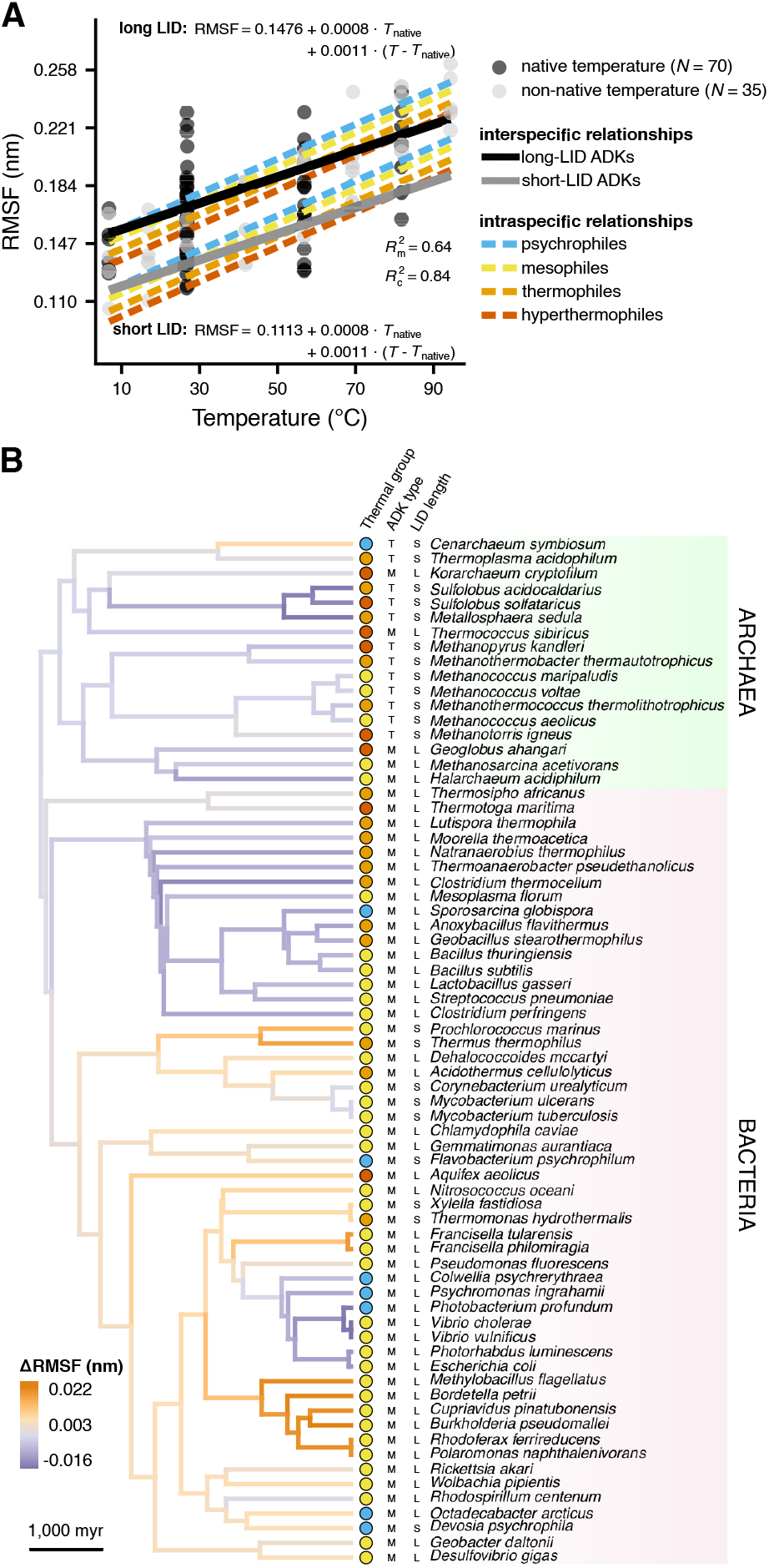
The relationship between RMSF (a measure of structural flexibility) and temperature. (A) Flexibility increases with temperature, across and within species. Note that intraspecific relationships (dashed lines) are plotted separately for long-LID (top) and short-LID ADKs (bottom). (B) Distribution of the phylogenetic random effect on the intercept across the branches of the species’ phylogeny. Each branch is coloured based on its difference from the intercept of the corresponding relationship in panel A. Panel B also shows the thermal group for each species, the ADK type (M for monomeric, T for trimeric), and the LID length (S for short, L for long).

### Impacts of temperature on ADK compactness

Following a similar approach to that in the previous subsection, we detected a statistically supported albeit very weak intraspecific decline in SASA (a surface compactness proxy) with temperature (Fig. 6A). In contrast, a systematic interspecific relationship was not supported. Temperature explained a negligible amount of variance in SASA (1%), with the overwhelming majority (95%) being captured by the phylogenetic random effect on the intercept. We verified the low explanatory power of temperature for SASA by estimating the intraspecific relationship separately for each of the five species for which simulations were performed at multiple temperatures. This showed that SASA systematically decreased with temperature for only three out of five species, with this decrease being non-negligible for only two species (Supplementary Fig. S5). The two species with the steepest intraspecific slopes (*Aquifex aeolicus*, a hyperthermophile and *Sporosarcina globispora*, a psychrophile) lie on opposite ends of the temperature spectrum which prevents us from drawing any further mechanistic conclusions. Examining the phylogenetic distribution of the estimated random effect revealed that almost all bacterial lineages had evolved relatively high SASA values, in contrast to archaeal lineages, many of which had shifted towards lower SASA (Fig. 6B).

**Fig. 6:**
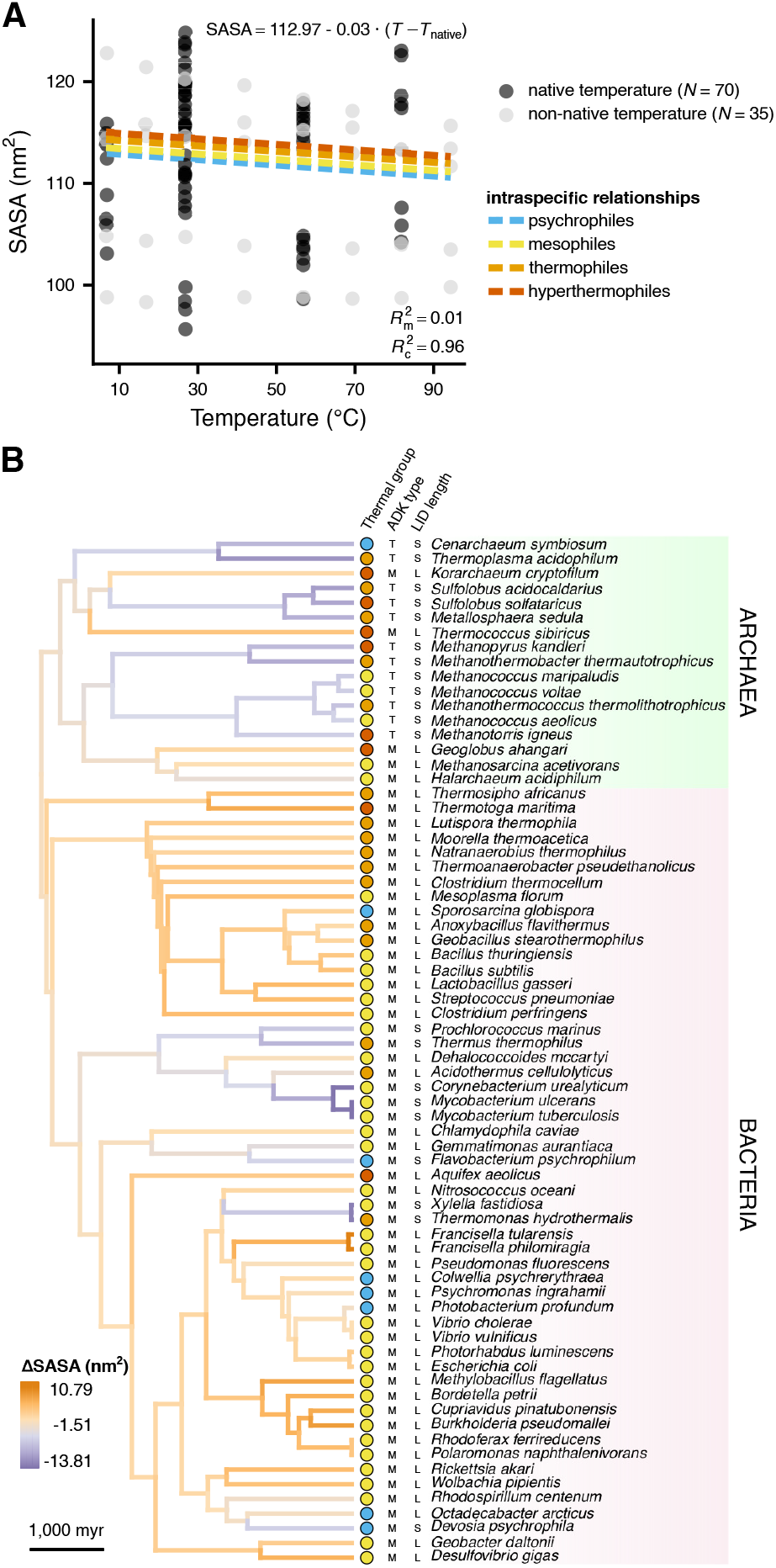
The relationship between solvent-accessible surface area (SASA) and temperature. (A) While SASA statistically decreases with temperature within species, this accounts for only 1% of the variance. (B) Distribution of the phylogenetic random effect on the intercept across the branches of the species’ phylogeny, as in Fig. 5.

A systematic relationship between temperature and r_gyr_ (a measure of overall structure compactness), could not be found, either within or across species (Fig. 7 and Supplementary Fig. S6). Long- and short-LID ADKs differed in r_gyr_ by 0.1 nm on average, which explained 31% of variance. 55% of the remaining variance was captured by a phylogenetic random effect on the intercept. These results indicate that neither acute nor long-term exposure to elevated temperatures systematically influence the overall compactness of prokaryotic ADKs.

**Fig. 7:**
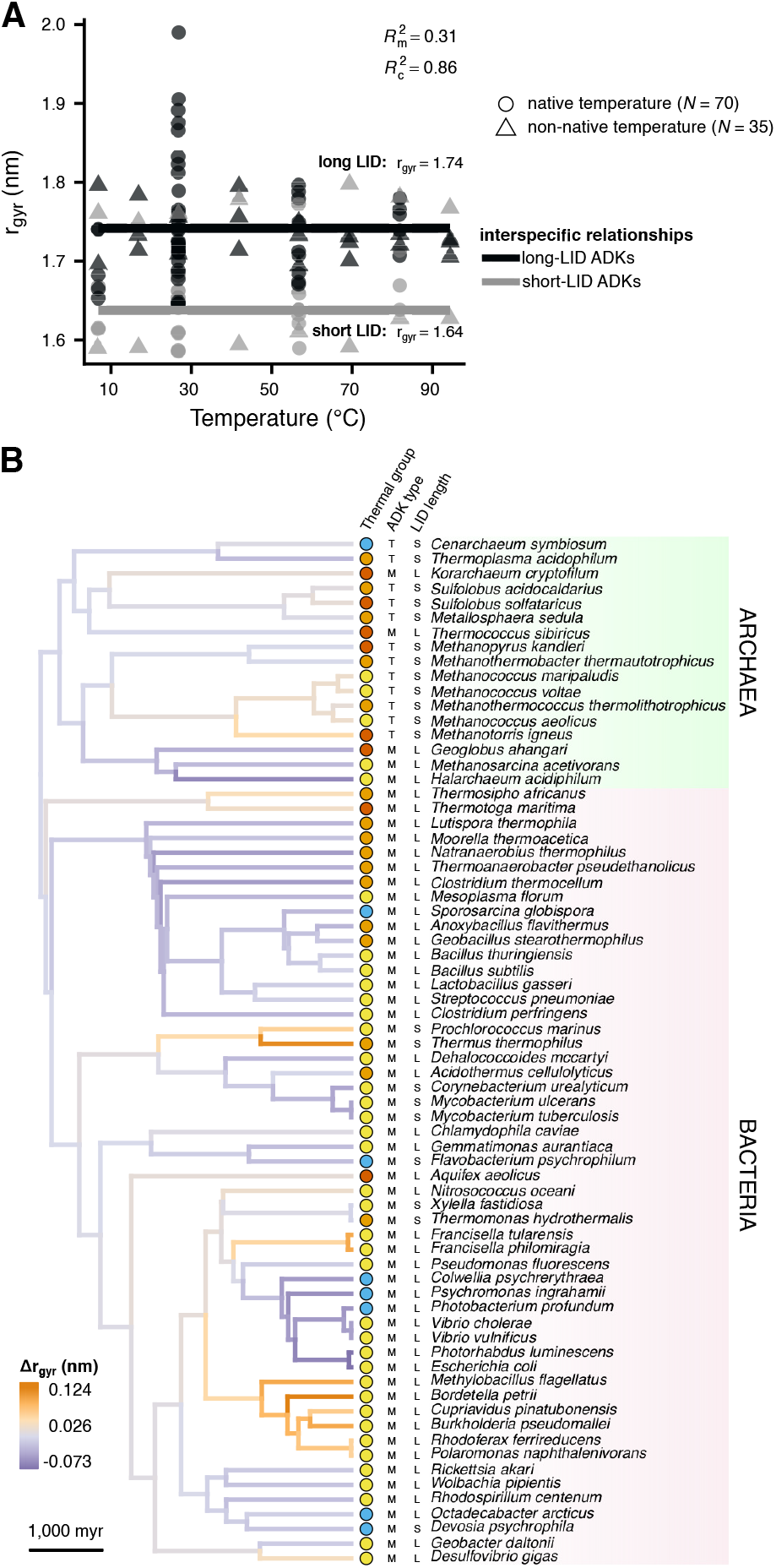
The relationship between the radius of gyration (a measure of overall structure compactness) and temperature. (A) Temperature does not systematically affect the radius of gyration, either within or across species. Long- and short-LID ADKs have slightly different intercepts. (B) Distribution of the phylogenetic random effect on the intercept across the branches of the species’ phylogeny, as in Fig. 5.

### Amino acid contacts and thermal adaptation

Our third key question was whether contacts among amino acids play a major role in the adaptation of ADKs to different thermal environments. We addressed this question in two ways. First, we examined if there is a systematic interspecific relationship between contact counts and temperature, separately for each contact type (see Fig. 4). We found that adaptation to higher temperatures is weakly associated with a small increase in the number of salt bridges and slight declines in negatively charged - polar and polar - polar contact counts. Specifically, compared to psychrophilic ADKs, hyperthermophilic orthologs have on average three more salt bridges, two fewer negatively charged - polar contacts, and three fewer polar - polar contacts (Fig. 8). In contrast, the number of positively charged - polar contacts was independent of temperature. It is worth pointing out here that these relationships were quite noisy, with temperature explaining between 22% and 51% of variance.

**Fig. 8:**
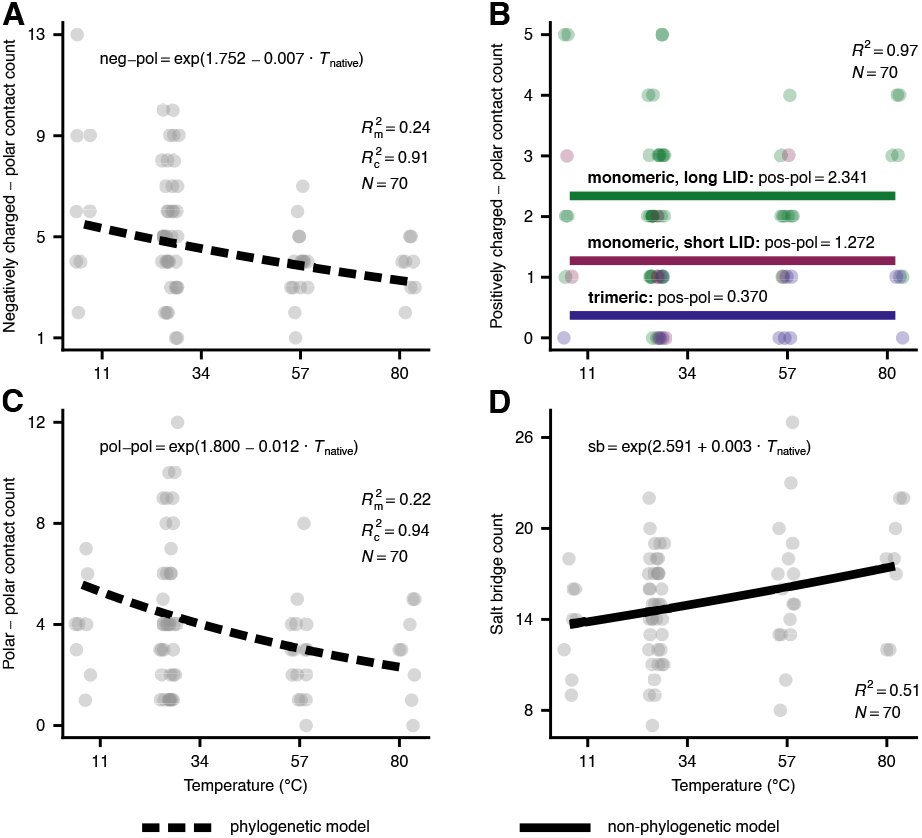
The impact of temperature on the counts of the four contact types. Each data point corresponds to a different ADK, whereas lines stand for the best-fitting model per panel (see Methods). In panel B, different colors are used for trimeric ADKs, monomeric ADKs with a short LID, and monomeric ADKs with a long LID. To reduce the extensive overlap among data points (especially in panel B), we have slightly displaced data points along the horizontal axis. See also Supplementary Figs. S3 and S4 for the distributions of the phylogenetic random effects of the models shown in panels A and C.

Second, we investigated whether thermal adaptation may be linked to the presence or absence of several key contacts at specific regions of the protein. For the latter, we inferred the average contact network for the monomeric ADKs of each thermal group. More precisely, for all possible pairs of alignment columns, we estimated the probability of the presence of a contact while accounting for the evolutionary relationships among species. We then built networks that included all contacts with a probability of at least 0.4 (Fig. 9). Overall, we detected significant within-group variation in the presence and absence of amino acid contacts, with very few contacts having probabilities close to 1 within a given thermal group (see also Supplementary Fig. S7 where we raised the probability cutoff from 0.4 to 0.7). It is worth noting that *∼* 93.5% of all observed contacts had a probability below 0.4 and, hence, are not shown in the average networks of Fig. 9. Furthermore, most of the commonly occurring contacts were either shared by all four thermal groups (e.g., salt bridges in the LID domain) or only shared by non-adjacent thermal groups (e.g., the positively charged - polar contact between alignment columns 77 and 83 of psychrophiles and thermophiles).

**Fig. 9:**
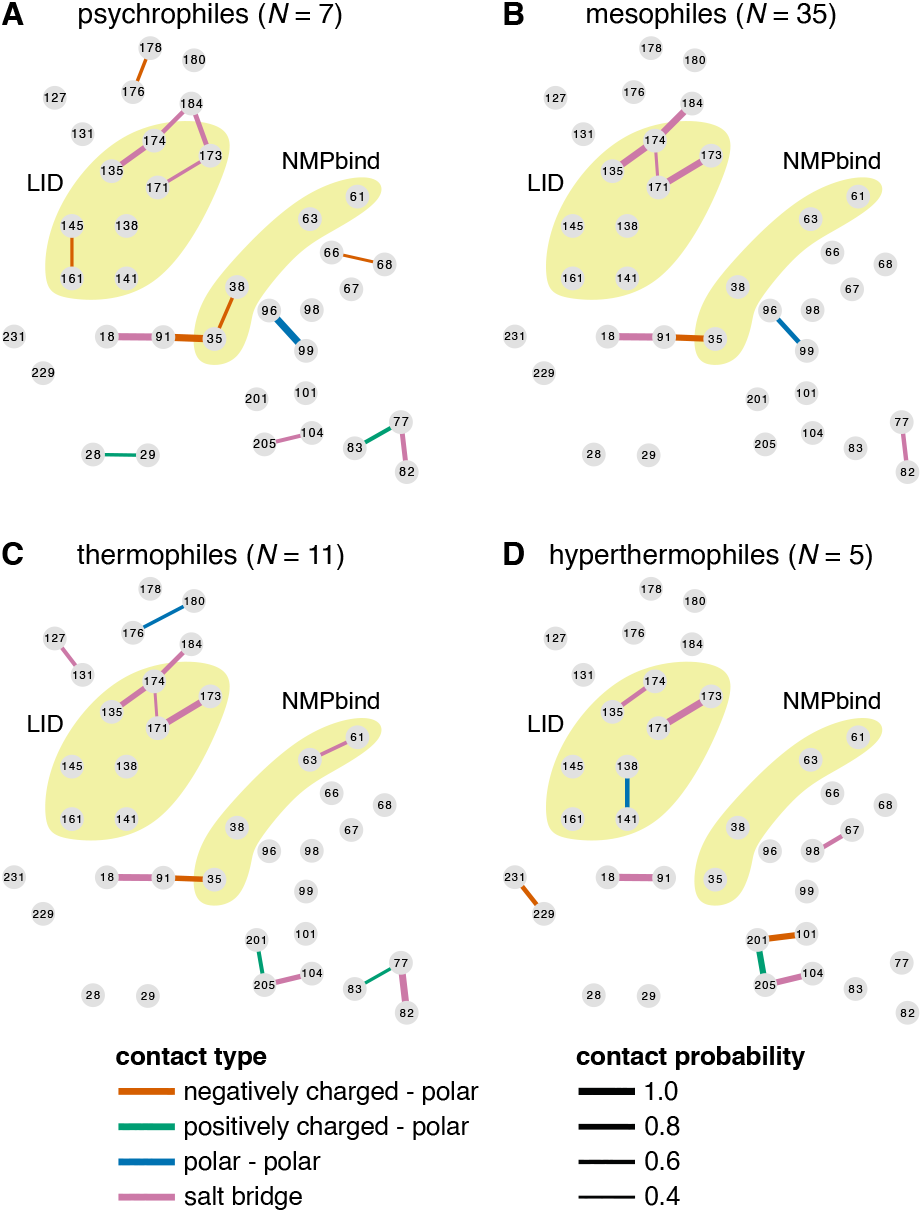
Average contact networks for each thermal group. Each node corresponds to a different column of the amino acid alignment of monomeric ADKs. Nodes are positioned approximately where they would lie along the ADK structure (see Fig. 2), with the LID and NMPbind domains shown in yellow. The four contact types are also shown in different colours, with line thickness being proportional to the probability of each contact. Contacts with a probability below 0.4 are not shown.

These findings indicate that adaptation of ADKs to extreme low- or high-temperature environments is not tightly linked to the presence or absence of specific contacts among amino acid sidechains. Instead, different lineages appear to respond to similar thermal challenges with very different evolutionary solutions.

## Concluding remarks

Our study shows that thermal adaptation of prokaryotic adenylate kinases is achieved at the structural level in diverse ways (Fig. 4). In particular, we found evidence for a systematic increase in global structural flexibility with temperature, both within species (as expected; see Fig. 1), but also across species (Fig. 5). This means that psychrophilic orthologs are on average a) more flexible than their warmer-adapted orthologs at any given temperature and b) less flexible than their warmer-adapted orthologs when compared at their respective native temperature ranges. The aforementioned pattern does not support the “corresponding states” hypothesis (Fig. 1), which would predict an interspecific slope of zero. In contrast to structural flexibility, a change in structural compactness, either only at the solvent-exposed surface area of the protein or overall, could not be systematically linked to thermal adaptation (Figs. 6 and 7). Similarly, the topology and types of contacts formed among pairs of amino acids varied considerably across adenylate kinases from similar thermal environments (Figs. 8 and 9). Overall, these results are consistent with the presence of numerous lineage-specific mechanisms for thermal adaptation that may in part reflect the thermal conditions experienced by the ancestors of each lineage (Berezovsky and Shakhnovich, 2005).

A limitation of our study is that we did not include any kinetic variables in our analyses, i.e., *k*_cat_ (turnover number; the catalytic velocity of a single enzyme under saturating substrate conditions), *K*_M_ (the Michaelis constant; a measure of substrate binding affinity), and their ratio (a measure of catalytic efficiency). Such variables would enable us to test for a trade-off between structural stability and catalytic activity, as has been previously suggested (e.g., Miller 2017; Nguyen et al. 2017; Maffucci et al. 2020; Stark et al. 2022). Unfortunately, published estimates of these parameters for prokaryotic ADKs had been almost completely nonexistent. Nevertheless, a recent study by Muir et al. (2024) produced a dataset of kinetic parameters for 193 prokaryotic ADKs, 17 of which were part of our dataset. Across those 17 ADKs, there was no systematic relationship between temperature-normalised flexibility and kinetic parameters (Supplementary Section S6). This pattern suggests the absence of a stability-activity trade-off in prokaryotic ADKs, in line with the conclusions of Muir et al. (2024). Given the importance of ADK for cellular energy homeostasis and, hence, fitness (Couñago and Shamoo, 2005; Couñago et al., 2006, 2008; Peña et al., 2010; Tükenmez et al., 2016), the lack of this trade-off may arise from strong selection for sufficient ADK activity in diverse thermal environments. In any case, a possible link between the strength of the stability-activity trade-off and the importance of a given enzyme for fitness remains to be explored in future studies.

It is worth pointing out that adaptation of enzyme structures is but one way through which species respond to thermal challenges. Other mechanisms include a) evolving multiple gene transcripts or gene copies with different thermal characteristics (Sanchez-Perez et al., 2008; Telonis-Scott et al., 2014), b) tweaking the expression levels of heat shock proteins (Oksala et al., 2014), c) changes in metabolic networks (Braakman et al., 2017), d) modifications in the fluidity of cellular membranes (Ernst et al., 2016), and e) behavioural responses for locating more thermally suitable (micro-)habitats (Freitas et al., 2015; Ramalho et al., 2023). Such mechanisms are not mutually exclusive, with their relative importance varying across species, depending on their physiological, morphological, or other traits and the characteristics of their local environment.

Overall, our study provides quantitative estimates of the intra- and interspecific relationships between environmental temperature and several structural variables of prokaryotic adenylate kinases, as well as any phylogenetically-structured deviations from such relationships (e.g., see Fig. 5B). Future studies could focus on obtaining similar estimates for many other enzymes from diverse taxonomic groups, which would ultimately pave the way for systematic, genome-wide comparisons of the relative importance of different structural modifications for the thermal adaptation of enzymes.

## Methods

### Compilation of the dataset of ADK structures

The ADKs included in the present study were chosen between October 2016 and February 2018. At that time, there were fewer than 20 species of bacteria and archaea with an experimentally determined ADK structure available in the Protein Data Bank. Thus, to obtain a larger and more diverse dataset, we manually selected further ADK sequences from the UniProt database (The UniProt Consortium, 2017), representing a wide range of clades and thermal environments. To estimate the structure of each ADK sequence, we performed homology modelling with the Phyre2 server (Kelley et al., 2015). This approach uses structural information from one or more experimentally determined ADKs (templates) to produce structural models for ADKs for which only the sequence is available (queries). More specifically, we used the intensive mode of the Phyre2 server which automatically selects the most appropriate template(s) for each query sequence, based on the confidence and coverage of the sequence alignment between query and template(s). This method also incorporates an *ab initio* step to model amino acids that are not covered by the template(s). To ensure that the quality of all resulting models was sufficiently high, we ensured that all 70 ADKs had no more than four non-covered amino acids and performed additional quality evaluation tests (Supplementary Fig. S1).

### Estimation of evolutionary trees

#### Species tree

To obtain a time-calibrated phylogeny for the species in our ADK dataset, we extracted a tree topology—that included all 70 species—from the Open Tree of Life (OTL; Hinchliff et al. 2015), and a time-calibrated phylogeny—for a subset of them— from the TimeTree database (Kumar et al., 2022). We then used the congruify.phylo function of the geiger R package (v. 2.0.11; Pennell et al. 2014) to transfer age estimates from the TimeTree phylogeny to that of the OTL, where possible (Eastman et al., 2013). The remaining nodes were time-calibrated using the treePL software (Smith and O’Meara, 2012).

#### ADK gene tree

To reconstruct the ADK gene tree, we first aligned the 70 amino acid sequences with MAFFT (v. 7.490; Katoh and Standley 2013) using the G-INS-i global alignment algorithm. We next ran IQ-TREE (v. 2.3.5; Minh et al. 2020) to identify the best-fitting amino acid substitution model, and perform 300 maximum-likelihood tree searches and 300 non-parametric bootstrap replicates. Finally, we mapped bootstrap support values to the maximum-likelihood tree with the highest log-likelihood.

### Molecular dynamics simulations

#### Simulation protocol

Molecular dynamics simulations were conducted with the Desmond software suite and its graphical interface, Maestro (Version 10.6.013, Release 2016-2). Specifically, for each ADK structure, we first capped the N- and C-termini, added missing hydrogen atoms, and performed energy minimization in vacuum under the OPLS 2005 force field (Banks et al., 2005) to remove potential atom clashes. We then solvated the resulting structure in a periodic orthorhombic simulation box with Simple Point Charge water molecules (Berendsen et al., 1981), and with Na^+^ and Cl^*−*^ ions added for neutralizing the charge of the protein and for generating a salt concentration of 0.15 M. Box volumes ranged between *∼*207 and *∼*360 nm^3^, with the median being at *∼*248 nm^3^. Each system was next relaxed through Desmond’s standard protocol, following which we performed ten replicate simulations of 200 ns per ADK. In these simulations, the temperature was held constant using the Nosé-Hoover thermostat (Martyna et al., 1992) at a different value per thermal group (see the “Comparison of the structural parameter space occupied by the four thermal groups” subsection above). Similarly, we ensured that the pressure of the system was held constant at the default value of 1.01325 bar by enforcing isotropic coupling and a relaxation time of 2 ps using the Martyna-Tobias-Klein barostat (Martyna et al., 1994). For van der Waals and short-range electrostatic interactions, we set the cutoff radius to 0.9 nm. Finally, we set the integration time step to 2 fs (the default value) and recorded a snapshot of the system every 0.5 ns.

We also performed simulations at 7 additional temperatures (from 6.85^*°*^C to 94.35^*°*^C) for the ADKs of a) *Sporosarcina globispora* (a psychrophilic bacterium), b) *Mycobacterium tuberculosis* (a mesophilic bacterium), c) *Methanosarcina acetivorans* (a mesophilic archaeon), d) *Thermus thermophilus* (a thermophilic bacterium), and e) *Aquifex aeolicus* (a hyperthermophilic bacterium)

All molecular dynamics simulations were executed on desktop machines equipped with CUDA-enabled GPUs (e.g., GeForce GTX TITAN, Tesla P100) for a total time of *∼*35,300 hours. We would like to note that this runtime is a small fraction of what we would expect if we had performed the simulations on non-GPU systems.

#### Analysis of simulation trajectories

We used the Gromacs 2023 package (Abraham et al., 2015) to calculate the root mean square fluctuation (RMSF) of alpha carbon atoms, the solvent-accessible surface area (SASA), and the radius of gyration of alpha carbon atoms (*r*_gyr_). To measure the number of contacts among amino acid side chains, we used the MDAnalysis Python package (Michaud-Agrawal et al., 2011; Gowers et al., 2016). Specifically, we applied a typically-used distance cutoff of 0.5 nm between the centres of the interacting chemical groups of the two amino acids (e.g., carboxyl group in Asp and Glu, or guanidine group in Arg). From the resulting contacts, we excluded those that occurred for less than 50% of simulation time. Finally, we classified contacts based on the properties of the participating amino acids. Specifically, we treated a) Arg and Lys as positively charged, b) Asp and Glu as negatively charged, and c) Asn, Gln, Ser, Thr, and Tyr as polar amino acids.

### Phylogenetic comparative analyses

#### Phylogenetic PCA

We examined the distribution of the 70 ADKs throughout the structural parameter space by applying a phylogenetic PCA (Revell, 2009) to the matrix of structural variables, estimated at one (native) temperature per ADK. More precisely, we used the phyl.pca function of the phytools R package (v.2.1-1; Revell 2024), accounting for the level of phylogenetic signal in the data by simultaneously optimizing the *λ* parameter. Given that there was not adequate phylogenetic signal for inferring a statistically robust ADK gene tree (Fig. 3E), we instead used the species’ phylogeny as a proxy of the evolutionary relationships among ADKs.

#### Estimation of any intra- and interspecific relationships of RMSF, SASA, and r_gyr_ with temperature

To understand if RMSF, SASA, and r_gyr_ vary systematically with temperature within or across species, we fitted 32 alternative generalised linear mixed models for each of these three variables with the MCMCglmm R package (v. 2.36; Hadfield 2010). RMSF, SASA, and and r_gyr_ were treated as Gaussian-distributed response variables, with the measurement uncertainty of each data point explicitly accounted for, making each model “meta-analytic”. Possible explanatory variables were a) the native temperature (for the interspecific slope), b) the difference between the non-native and native temperature (if applicable, for the intraspecific slope), c) the ADK type (monomeric or trimeric), and d) the length of the LID (short or long). We fitted models with all possible combinations of these explanatory variables, as well as an empty (intercept-only) model. We also specified a phylogenetic variant of each model by adding a phylogenetic random effect on the intercept. The priors that we used were relatively uninformative, namely the default Gaussian prior for fixed effects, a parameter-expanded prior for the phylogenetic random effect (if applicable), and an inverse Gamma prior for the residual variance. For each model, we executed 3 independent chains for 1 million generations, with posterior samples obtained every 100 generations after discarding the first 100,000 as burn-in. We ensured that the 3 chains per model had converged to statistically equivalent posterior distributions and that the latter were sufficiently sampled by verifying that, for each model parameter, i) the effective sample size was at least 1,000 and ii) the potential scale reduction factor was below 1.1. To identify the best-fitting model for each response variable, we first excluded models if any of their fixed-effects coefficients (other than the intercept) had a 95% Highest Posterior Density interval that included zero. From the remaining models, we selected the one with the lowest Deviance Information Criterion value (Spiegelhalter et al., 2002). For this model, we calculated the marginal and conditional *R*^2^ values following Hadfield and Nakagawa (2010). 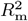 stands for the proportion of variance that is captured by the fixed effects, whereas 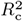 stands for the variance captured by both the fixed and the random effects.

#### Linking contacts to thermal adaptation

To test for any interspecific relationships between contact counts and temperature, we followed the approach described in the previous subsection. The main differences were that the response variables were treated as Poisson-distributed and that we did not attempt to estimate any intraspecific relationships. For the 3 independent chains to reach convergence, we set the number of MCMC generations to 50 million, the burn-in to 5 million, and sampled from the posterior every 15,000 generations.

For the contact networks analysis, we used the amino acid alignment of the 58 monomeric ADKs to objectively compare the observed contacts across ADKs. More precisely, we created a unique identifier for each contact, based on the alignment column numbers of the two participating amino acids. We next constructed matrices where the rows were the 58 ADKs and the columns were the contacts. We made separate matrices for each thermal group and each contact type. The values in these matrices were either 0 (the contact is missing) or 1 (the contact is present). For columns where all values were 0 or 1, the contact probability is necessarily equal to 0 or 1, respectively. For all remaining matrix columns, we fitted a phylogenetic threshold model (Felsenstein, 2005; Hadfield, 2015) with MCMCglmm, separately for each thermal group. This type of model assumes that the discrete response variable (the presence/absence vector of a given contact in this case) is governed by an unobserved continuous variable, called the “liability”. The contact should be absent if the liability is below 0 and present otherwise. To this end, each threshold model had the presence/absence vector as the response variable, no predictors other than an intercept, and a phylogenetic random effect on the intercept. The priors were set similarly to those in the previous subsection, except for the residual variance which is not identifiable in threshold models and had to be fixed to 1. We executed the 3 chains per model for 500,000 generations, removed samples from the first 50,000 generations as burn-in, and then sampled from the posterior every 50 generations. After running convergence and sampling diagnostics as previously described, we calculated the contact probability based on the posterior distribution of the intercept. Because the intercept stands for the phylogenetically corrected liability for a given contact and thermal group, the contact probability can be calculated as the number of posterior samples of the intercept with a value of 0 or higher, divided by the total number of samples.

## Data and code availability

The data underlying this article are available from Figshare at https://doi.org/10.6084/m9.figshare.28436891 [dataset] Kontopoulos et al. (2025). The source code for the analyses of the present study is available from Codeberg at https://codeberg.org/dgkontopoulos/Kontopoulos_et_al_evolution_of_ADK_structures_2025 and archived on Zenodo at https://doi.org/10.5281/zenodo.14891812 [code] Kontopoulos et al. (2025).

## Author contributions

DGK conceived the study, designed the methodology, collected and analysed data, and created figures. IP provided guidance on the protocol for the molecular dynamics simulations and extracted contact counts from simulation trajectories. TGB and SP supervised the study and contributed to its methodology. DGK wrote the initial draft, which was later revised based on inputs from all authors.

## Acknowledgements

We are grateful to Imperial College London’s Research Computing Service (DOI: 10.14469/hpc/2232) for providing the necessary computational resources for the molecular dynamics simulations described in this study.

## References

Abraham, M. J., T. Murtola, R. Schulz, S. Páll, J. C. Smith, B. Hess, and E. Lindahl. 2015. GROMACS: High performance molecular simulations through multi-level parallelism from laptops to supercomputers. SoftwareX 1:19–25.

Banks, J. L., H. S. Beard, Y. Cao, A. E. Cho, W. Damm, R. Farid, A. K. Felts, T. A. Halgren, D. T. Mainz, J. R. Maple, et al. 2005. Integrated modeling program, applied chemical theory (IMPACT). Journal of Computational Chemistry 26:1752–1780.

Bellinzoni, M., A. Haouz, M. Graña, H. Munier-Lehmann, W. Shepard, and P. M. Alzari. 2006. The crystal structure of Mycobacterium tuberculosis adenylate kinase in complex with two molecules of ADP and Mg2+ supports an associative mechanism for phosphoryl transfer. Protein Science 15:1489– 1493.

Berendsen, H. J., J. P. Postma, W. F. van Gunsteren, and J. Hermans. 1981. Interaction models for water in relation to protein hydration. in Intermolecular forces. Pp. 331–342. Springer.

Berezovsky, I. N., and E. I. Shakhnovich. 2005. Physics and evolution of thermophilic adaptation. Proceedings of the National Academy of Sciences 102:12742–12747.

Braakman, R., M. J. Follows, and S. W. Chisholm. 2017. Metabolic evolution and the self-organization of ecosystems. Proceedings of the National Academy of Sciences 114:E3091– E3100.

Cipolla, A., F. Delbrassine, J.-L. Da Lage, and G. Feller. 2012. Temperature adaptations in psychrophilic, mesophilic and thermophilic chloride-dependent alpha-amylases. Biochimie 94:1943–1950.

Clarke, A. 2017. Principles of Thermal Ecology: Temperature, Energy, and Life. Oxford University Press.

Collins, T., M.-A. Meuwis, C. Gerday, and G. Feller. 2003. Activity, stability and flexibility in glycosidases adapted to extreme thermal environments. Journal of Molecular Biology 328:419–428.

Couñago, R., S. Chen, and Y. Shamoo. 2006. In vivo molecular evolution reveals biophysical origins of organismal fitness. Molecular Cell 22:441–449.

Couñago, R., and Y. Shamoo. 2005. Gene replacement of adenylate kinase in the gram-positive thermophile Geobacillus stearothermophilus disrupts adenine nucleotide homeostasis and reduces cell viability. Extremophiles 9:135–144.

Couñago, R., C. J. Wilson, M. I. Pena, P. Wittung-Stafshede, and Y. Shamoo. 2008. An adaptive mutation in adenylate kinase that increases organismal fitness is linked to stability–activity trade-offs. Protein Engineering, Design & Selection 21:19–27.

Diessner, E. M., G. R. Takahashi, C. T. Butts, and R. W. Martin. 2024. Comparative analysis of thermal adaptations of extremophilic prolyl oligopeptidases. Biophysical Journal.

Eastman, J. M., L. J. Harmon, and D. C. Tank. 2013. Congruification: support for time scaling large phylogenetic trees. Methods in Ecology and Evolution 4:688–691.

Engqvist, M. K. 2018. Correlating enzyme annotations with a large set of microbial growth temperatures reveals metabolic adaptations to growth at diverse temperatures. BMC Microbiology 18:1–14.

Ernst, R., C. S. Ejsing, and B. Antonny. 2016. Homeoviscous adaptation and the regulation of membrane lipids. Journal of Molecular Biology 428:4776–4791.

Feil, E., G. Carpenter, and B. G. Spratt. 1995. Electrophoretic variation in adenylate kinase of Neisseria meningitidis is due to inter-and intraspecies recombination. Proceedings of the National Academy of Sciences 92:10535–10539.

Feil, E., J. Zhou, J. M. Smith, and B. G. Spratt. 1996. A comparison of the nucleotide sequences of the adk and recA genes of pathogenic and commensal Neisseria species: evidence for extensive interspecies recombination within adk. Journal of Molecular Evolution 43:631–640.

Feller, G. 2010. Protein stability and enzyme activity at extreme biological temperatures. Journal of Physics: Condensed Matter 22:323101.

Felsenstein, J. 2005. Using the quantitative genetic threshold model for inferences between and within species. Philosophical Transactions of the Royal Society B: Biological Sciences 360:1427–1434.

Fitter, J., R. Herrmann, N. Dencher, A. Blume, and T. Hauss. 2001. Activity and stability of a thermostable α-amylase compared to its mesophilic homologue: mechanisms of thermal adaptation. Biochemistry 40:10723–10731.

Freitas, C., E. M. Olsen, E. Moland, L. Ciannelli, and H. Knutsen. 2015. Behavioral responses of atlantic cod to sea temperature changes. Ecology and Evolution 5:2070–2083.

Gowers, R. J., M. Linke, J. Barnoud, T. J. E. Reddy, M. N. Melo, S. L. Seyler, J. Domański D. L. Dotson, S. Buchoux, I. M. Kenney, and O. Beckstein. 2016. MDAnalysis: A Python Package for the Rapid Analysis of Molecular Dynamics Simulations. in Sebastian Benthall and Scott Rostrup, eds. Proceedings of the 15th Python in Science Conference. Pp. 98–105.

Hadfield, J., and S. Nakagawa. 2010. General quantitative genetic methods for comparative biology: phylogenies, taxonomies and multi-trait models for continuous and categorical characters. Journal of Evolutionary Biology 23:494–508.

Hadfield, J. D. 2010. MCMC methods for multi-response generalized linear mixed models: the MCMCglmm R package. Journal of Statistical Software 33:1–22.

Hadfield, J. D.. 2015. Increasing the efficiency of MCMC for hierarchical phylogenetic models of categorical traits using reduced mixed models. Methods in Ecology and Evolution 6:706–714.

Hajdú, I., C. Bőthe, A. Szilágyi, J. Kardos, P. Gál, and P. Závodszky. 2008. Adjustment of conformational flexibility of glyceraldehyde-3-phosphate dehydrogenase as a means of thermal adaptation and allosteric regulation. European Biophysics Journal 37:1139–1144.

Hinchliff, C. E., S. A. Smith, J. F. Allman, J. G. Burleigh, R. Chaudhary, L. M. Coghill, K. A. Crandall, J. Deng, B. T. Drew, R. Gazis, et al. 2015. Synthesis of phylogeny and taxonomy into a comprehensive tree of life. Proceedings of the National Academy of Sciences 112:12764–12769.

Jenney Jr, F. E., H. Wang, S. J. George, J. Xiong, Y. Guo, L. B. Gee, J. J. Marizcurrena, S. Castro-Sowinski, A. Staskiewicz, Y. Yoda, M. Y. Hu, K. Tamasaku, N. Nagasawa, L. Li, H. Matsuura, T. Doukov, and S. P. Cramer. 2024. Temperature-dependent iron motion in extremophile rubredoxins–no need for ‘corresponding states’. Scientific Reports 14:12197.

Karshikoff, A., and R. Ladenstein. 1998. Proteins from thermophilic and mesophilic organisms essentially do not differ in packing. Protein Engineering 11:867–872.

Katoh, K., and D. M. Standley. 2013. MAFFT multiple sequence alignment software version 7: improvements in performance and usability. Molecular Biology and Evolution 30:772–780.

Kelley, L. A., S. Mezulis, C. M. Yates, M. N. Wass, and M. J. Sternberg. 2015. The Phyre2 web portal for protein modeling, prediction and analysis. Nature Protocols 10:845–858.

Knapp, B. D., and K. C. Huang. 2022. The effects of temperature on cellular physiology. Annual Review of Biophysics 51:499–526.

[code] Kontopoulos, D.-G., I. Patmanidis, T. G. Barraclough, and S. Pawar. 2025. Code from: Changes in flexibility but not in compactness underlie the thermal adaptation of prokaryotic adenylate kinases. dgkontopoulos/konto poulos et al evolution of adk structures 2025. Zenodo URL 10.5281/zenodo.14891813.

[dataset] Kontopoulos, D.-G., I. Patmanidis, T. G. Barraclough, and S. Pawar. 2025. Data from: Changes in flexibility but not in compactness underlie the thermal adaptation of prokaryotic adenylate kinases. Figshare URL 10.6084/m9.figshare.28436891.

Kovermann, M., C. Grundström, A. E. Sauer-Eriksson, U. H. Sauer, and M. Wolf-Watz. 2017. Structural basis for ligand binding to an enzyme by a conformational selection pathway. Proceedings of the National Academy of Sciences 114:6298– 6303.

Kumar, S., M. Suleski, J. M. Craig, A. E. Kasprowicz,M. Sanderford, M. Li, G. Stecher, and S. B. Hedges. 2022. TimeTree 5: an expanded resource for species divergence times. Molecular Biology and Evolution 39:msac174.

Li, D., M. S. Liu, and B. Ji. 2015. Mapping the dynamics landscape of conformational transitions in enzyme: the adenylate kinase case. Biophysical Journal 109:647–660.

Livingstone, C. D., and G. J. Barton. 1993. Protein sequence alignments: a strategy for the hierarchical analysis of residue conservation. Bioinformatics 9:745–756.

Maffucci, I., D. Laage, G. Stirnemann, and F. Sterpone. 2020. Differences in thermal structural changes and melting between mesophilic and thermophilic dihydrofolate reductase enzymes. Physical Chemistry Chemical Physics 22:18361–18373.

Martyna, G. J., M. L. Klein, and M. Tuckerman. 1992. Nosé–Hoover chains: The canonical ensemble via continuous dynamics. The Journal of Chemical Physics 97:2635–2643.

Martyna, G. J., D. J. Tobias, and M. L. Klein. 1994. Constant pressure molecular dynamics algorithms. The Journal of Chemical Physics 101:4177–4189.

Michaud-Agrawal, N., E. J. Denning, T. B. Woolf, and O. Beckstein. 2011. MDAnalysis: a toolkit for the analysis of molecular dynamics simulations. Journal of Computational Chemistry 32:2319–2327.

Miller, S. R. 2017. An appraisal of the enzyme stability-activity trade-off. Evolution 71:1876–1887.

Minh, B. Q., H. A. Schmidt, O. Chernomor, D. Schrempf, M. D. Woodhams, A. Von Haeseler, and R. Lanfear. 2020. IQ-TREE 2: new models and efficient methods for phylogenetic inference in the genomic era. Molecular Biology and Evolution 37:1530– 1534.

Muir, D. F., G. P. R. Asper, P. Notin, J. A. Posner, D. S. Marks, M. J. Keiser, and M. M. Pinney. 2024. Evolutionary-scale enzymology enables biochemical constant prediction across a multi-peaked catalytic landscape. bioRxiv 2024.10.23.619915.

Nguyen, V., C. Wilson, M. Hoemberger, J. B. Stiller, R. V. Agafonov, S. Kutter, J. English, D. L. Theobald, and D. Kern. 2017. Evolutionary drivers of thermoadaptation in enzyme catalysis. Science 355:289–294.

Oksala, N. K., F. G. Ekmekçi, E. Özsoy, Ş. Kirankaya, T. Kokkola, G. Emecen, J. Lappalainen, K. Kaarniranta, and M. Atalay. 2014. Natural thermal adaptation increases heat shock protein levels and decreases oxidative stress. Redox Biology 3:25–28.

Pawar, S., A. I. Dell, and V. M. Savage. 2015. From metabolic constraints on individuals to the dynamics of ecosystems. Pp. 3–36. in Aquatic functional biodiversity. Elsevier.

Peña, M. I., E. Van Itallie, M. R. Bennett, and Y. Shamoo. 2010. Evolution of a single gene highlights the complexity underlying molecular descriptions of fitness. Chaos: An Interdisciplinary Journal of Nonlinear Science 20.

Pennell, M. W., J. M. Eastman, G. J. Slater, J. W. Brown, J. C. Uyeda, R. G. FitzJohn, M. E. Alfaro, and L. J. Harmon. 2014. geiger v2.0: an expanded suite of methods for fitting macroevolutionary models to phylogenetic trees. Bioinformatics 30:2216–2218.

Pettersen, E. F., T. D. Goddard, C. C. Huang, E. C. Meng, G. S. Couch, T. I. Croll, J. H. Morris, and T. E. Ferrin. 2021. UCSF ChimeraX: Structure visualization for researchers, educators, and developers. Protein Science 30:70–82.

Ramalho, Q., M. M. Vale, S. Manes, P. Diniz, A. Malecha, and J. A. Prevedello. 2023. Evidence of stronger range shift response to ongoing climate change by ectotherms and high-latitude species. Biological Conservation 279:109911.

Revell, L. J. 2009. Size-correction and principal components for interspecific comparative studies. Evolution 63:3258–3268.

Revell, L. J.. 2024. phytools 2.0: an updated R ecosystem for phylogenetic comparative methods (and other things). PeerJ 2:e16505.

Robinson-Rechavi, M., A. Alibés, and A. Godzik. 2006. ontribution of electrostatic interactions, compactness and quaternary structure to protein thermostability: lessons from ructural genomics of Thermotoga maritima. Journal of Molecular Biology 356:547–557.

chez-Perez, G., A. Mira, G. Nyirő, L. Pašić, and F. Rodriguez-alera. 2008. Adapting to environmental changes using pecialized paralogs. Trends in Genetics 24:154–158.

Smith, S. A., and B.C. O’Meara. 2012. treePL: divergence me estimation using penalized likelihood for large phylogenies. Bioinformatics 28:2689–2690.

Somero, G. N. 1978. Temperature adaptation of enzymes: biological optimization through structure-function ompromises. Annual Review of Ecology and Systematics 9:1–29.

Spiegelhalter, D. J., N. G. Best, B. P. Carlin, and A. Van Der Linde. 2002. Bayesian measures of model complexity and fit. Journal of the Royal Statistical Society: Series B (Statistical Methodology) 64:583–639.

Stark, C., T. Bautista-Leung, J. Siegfried, and D. Herschlag. 2022. Systematic investigation of the link between enzyme catalysis nd cold adaptation. eLife 11:e72884.

Szilagyi, A., and P. Závodszky. 2000. Structural differences etween mesophilic, moderately thermophilic and extremely thermophilic protein subunits: results of a comprehensive urvey. Structure 8:493–504.

Telonis-Scott, M., A. S. Clemson, T. K. Johnson, and C.M. Sgró. 2014. Spatial analysis of gene regulation reveals new insights into the molecular basis of upper thermal limits. Molecular cology 23:6135–6151.

The UniProt Consortium. 2017. UniProt: the universal protein nowledgebase. Nucleic Acids Research 45:D158–D169.

Tompa, D. R., M. M. Gromiha, and K. Saraboji. 2016. ontribution of main chain and side chain atoms and their locations to the stability of thermophilic proteins. Journal of olecular Graphics and Modelling 64:85–93.

Tukeenmez, H., H. M. Magnussen, M. Kovermann, A. Byström, nd M. Wolf-Watz. 2016. Linkage between fitness of yeast cells and adenylate kinase catalysis. PLoS One 11:e0163115.

Uyeda, J. C., R. Zenil-Ferguson, and M. W. Pennell. 2018. Rethinking phylogenetic comparative methods. Systematic iology 67:1091–1109.

Vázquez, D. P., E. Gianoli, W. F. Morris, and F. Bozinovic. 2017. Ecological and evolutionary impacts of changing climatic variability. Biological Reviews 92:22–42.

Vonrhein, C., H. Bönisch, G. Schäfer, and G. E. Schulz. 1998. The structure of a trimeric archaeal adenylate kinase. Journal f Molecular Biology 282:167–179.

Watterhouse, A. M., J. B. Procter, D. M. Martin, M. Clamp, and. J. Barton. 2009. Jalview Version 2—a multiple sequence alignment editor and analysis workbench. Bioinformatics 5:1189–1191.

Yu, G., D. K. Smith, H. Zhu, Y. Guan, and T. T.-Y. Lam. 2017. ggtree: an R package for visualization and annotation of phylogenetic trees with their covariates and other associated data. Methods in Ecology and Evolution 8:28–36.

Zhu, W., K. Sandman, G. E. Lee, J. N. Reeve, and M. F. Summers. 1998. NMR structure and comparison of the archaeal histone HFoB from the mesophile Methanobacterium formicicum with HMfB from the hyperthermophile Methanothermus fervidus. Biochemistry 37:10573–10580.

