## Supplementary Information for "Changes in flexibility but not in compactness underlie the thermal adaptation of prokaryotic adenylate kinases"

**S4 Species-specific relationships between temperature, RMSF, SASA, or $r_{gyr}$** 9

**S5 Average contact networks with a probability of at least 0.7** 11

**S6 Correlations between flexibility and kinetic parameters** 11

### S1 List of ADKs included in the present study

Table S1: Prokaryotic ADKs analysed and their main characteristics.

| Species | UniProt ID | Superkingdom | Thermal group | LID | Type |
|---|---|---|---|---|---|
| *Acidothermus cellulolyticus* | A0LRP1 | Bacteria | thermophilic | long | monomeric |
| *Anoxybacillus flavithermus* | B7GJ88 | Bacteria | thermophilic | long | monomeric |
| *Aquifex aeolicus* | O66490 | Bacteria | hyperthermophilic | long | monomeric |
| *Bacillus subtilis* | P16304 | Bacteria | mesophilic | long | monomeric |
| *Bacillus thuringiensis* | A0R8K1 | Bacteria | mesophilic | long | monomeric |
| *Bordetella petrii* | A9IQ39 | Bacteria | mesophilic | long | monomeric |
| *Burkholderia pseudomallei* | Q3JVB1 | Bacteria | mesophilic | long | monomeric |
| *Cenarchaeum symbiosum* | A0RUE3 | Archaea | psychrophilic | short | trimeric |
| *Chlamydophila caviae* | Q822Z1 | Bacteria | mesophilic | long | monomeric |
| *Clostridium perfringens* | Q0TMR7 | Bacteria | mesophilic | long | monomeric |
| *Clostridium thermocellum* | A3DJJ3 | Bacteria | thermophilic | long | monomeric |
| *Colwellia psychrerythraea* | Q47XA8 | Bacteria | psychrophilic | long | monomeric |
| *Corynebacterium urealyticum* | B1VEX6 | Bacteria | mesophilic | short | monomeric |
| *Cupriavidus pinatubonensis* | Q475G1 | Bacteria | mesophilic | long | monomeric |
| *Dehalococcoides mccartyi* | Q3Z960 | Bacteria | mesophilic | long | monomeric |
| *Desulfovibrio gigas* | C7U112 | Bacteria | mesophilic | long | monomeric |
| *Devosia psychrophila* | A0A0F5PRB6 | Bacteria | psychrophilic | short | monomeric |
| *Escherichia coli* | P69441 | Bacteria | mesophilic | long | monomeric |
| *Flavobacterium psychrophilum* | A6GZB9 | Bacteria | psychrophilic | short | monomeric |
| *Francisella philomiragia* | B0TXR2 | Bacteria | mesophilic | long | monomeric |
| *Francisella tularensis* | Q5NFR4 | Bacteria | mesophilic | long | monomeric |
| *Gemmatimonas aurantiaca* | C1A6S6 | Bacteria | mesophilic | long | monomeric |
| *Geobacillus stearothermophilus* | P27142 | Bacteria | thermophilic | long | monomeric |
| *Geobacter daltonii* | B9M6F7 | Bacteria | mesophilic | long | monomeric |
| *Geoglobus ahangari* | A0A0F7IDB9 | Archaea | hyperthermophilic | long | monomeric |
| *Halarchaeum acidiphilum* | U3AED7 | Archaea | mesophilic | long | monomeric |
| *Korarchaeum cryptofilum* | B1L775 | Archaea | hyperthermophilic | long | monomeric |
| *Lactobacillus gasseri* | Q046A5 | Bacteria | mesophilic | long | monomeric |
| *Lutispora thermophila* | A0A1M6DWR4 | Bacteria | thermophilic | long | monomeric |
| *Mesoplasma florum* | Q6F1X3 | Bacteria | mesophilic | long | monomeric |
| *Metallosphaera sedula* | A4YCZ0 | Archaea | thermophilic | short | trimeric |
| *Methanococcus aeolicus* | A6UW35 | Archaea | mesophilic | short | trimeric |
| *Methanococcus maripaludis* | A4FXE7 | Archaea | mesophilic | short | trimeric |
| *Methanococcus voltae* | P43411 | Archaea | mesophilic | short | trimeric |
| *Methanopyrus kandleri* | Q8TZB0 | Archaea | hyperthermophilic | short | trimeric |
| *Methanosarcina acetivorans* | Q8TRS3 | Archaea | mesophilic | long | monomeric |
| *Methanothermobacter thermautotrophicus* | O26135 | Archaea | thermophilic | short | trimeric |
| *Methanothermococcus thermolithotrophicus* | P43410 | Archaea | thermophilic | short | trimeric |
| *Methanotorris igneus* | P43408 | Archaea | hyperthermophilic | short | trimeric |
| *Methylobacillus flagellatus* | Q1GZB7 | Bacteria | mesophilic | long | monomeric |
| *Moorella thermoacetica* | Q2RFR8 | Bacteria | thermophilic | long | monomeric |
| *Mycobacterium tuberculosis* | P9WKF5 | Bacteria | mesophilic | short | monomeric |
| *Mycobacterium ulcerans* | A0PM98 | Bacteria | mesophilic | short | monomeric |
| *Natranaerobius thermophilus* | B2A4G2 | Bacteria | thermophilic | long | monomeric |
| *Nitrosococcus oceani* | Q3J7A3 | Bacteria | mesophilic | long | monomeric |
| *Octadecabacter arcticus* | M9RLZ6 | Bacteria | psychrophilic | long | monomeric |
| *Photobacterium profundum* | Q6LTE1 | Bacteria | psychrophilic | long | monomeric |
| *Photorhabdus luminescens* | Q7N0P5 | Bacteria | mesophilic | long | monomeric |
| *Polaromonas naphthalenivorans* | A1VNK2 | Bacteria | mesophilic | long | monomeric |
| *Prochlorococcus marinus* | A8G742 | Bacteria | mesophilic | short | monomeric |
| *Pseudomonas fluorescens* | C3KEC6 | Bacteria | mesophilic | long | monomeric |
| *Psychromonas ingrahamii* | A1STI3 | Bacteria | psychrophilic | long | monomeric |
| *Rhodoferax ferrireducens* | Q21TN6 | Bacteria | mesophilic | long | monomeric |
| *Rhodospirillum centenum* | B6IRS7 | Bacteria | mesophilic | long | monomeric |
| *Rickettsia akari* | A8GPC9 | Bacteria | mesophilic | long | monomeric |

Supplementary Table S1 – *Continued from the previous page*

| **Species** | **UniProt ID** | **Superkingdom** | **Thermal group** | | |
|---|---|---|---|---|---|
| *Sporosarcina globispora* | P84139 | Bacteria | psychrophilic | long | monomeric |
| *Streptococcus pneumoniae* | B8ZKQ1 | Bacteria | mesophilic | long | monomeric |
| *Sulfolobus acidocaldarius* | P35028 | Archaea | thermophilic | short | trimeric |
| *Sulfolobus solfataricus* | Q9UX83 | Archaea | hyperthermophilic | short | trimeric |
| *Thermoanaerobacter pseudethanolicus* | B0KCM0 | Bacteria | thermophilic | long | monomeric |
| *Thermococcus sibiricus* | C6A2Q1 | Archaea | hyperthermophilic | long | monomeric |
| *Thermomonas hydrothermalis* | A0A1M5AE38 | Bacteria | thermophilic | short | monomeric |
| *Thermoplasma acidophilum* | Q9HIT1 | Archaea | thermophilic | short | trimeric |
| *Thermosipho africanus* | B7IHW7 | Bacteria | thermophilic | long | monomeric |
| *Thermotoga maritima* | Q9X1I8 | Bacteria | hyperthermophilic | long | monomeric |
| *Thermus thermophilus* | Q5SHQ9 | Bacteria | thermophilic | short | monomeric |
| *Vibrio cholerae* | Q9KTB7 | Bacteria | mesophilic | long | monomeric |
| *Vibrio vulnificus* | Q7MMR5 | Bacteria | mesophilic | long | monomeric |
| *Wolbachia pipientis* | Q73HA6 | Bacteria | mesophilic | long | monomeric |
| *Xylella fastidiosa* | B2I757 | Bacteria | mesophilic | short | monomeric |

#### S2 Quality evaluation of structural models

We employed two criteria to evaluate the quality of our 70 ADK structures. First, we used the empirical equation of Rost (1999) to examine if the identity percentage between the query and template sequences was sufficiently high relative to the length of their alignment (Fig. S1A). This showed that all query-template alignments fall within the safe homology modelling zone of the parameter space and, therefore, the resulting models are expected to be reliable.

We also used the PROCHECK tool (Laskowski et al., 1993) to generate Ramachandran plots (Branden and Tooze, 2012) and examine the $\phi$ and $\psi$ backbone dihedral angles of amino acids in each ADK structure. The combinations of these angles can be classified—based on numerous experimentally determined structures from diverse proteins—as being part of a) disallowed regions, b) generously allowed regions, c) additional allowed regions, or d) the most favoured regions. In our case, all ADKs had at least 94.7% of their amino acids in their most favoured and additional allowed regions, whereas the proportion of amino acids in disallowed regions was never higher than 3.2% (Fig. S1B). These results further confirm the high quality of the structural models analysed in this study.

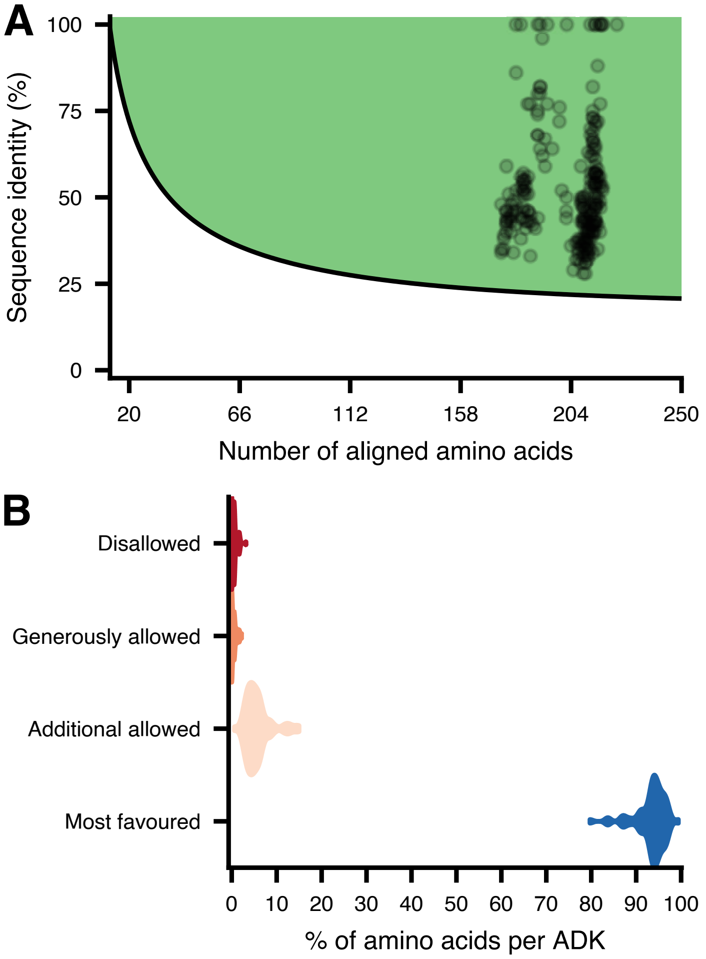

Figure S1: Tests for evaluation of the quality of ADK structures included in this study. (A) The relationship of the sequence identity between template and query sequences against the alignment length. The green area, defined by the empirical equation of Rost (1999), is expected to produce reliable homology models. Note that certain ADKs (those at 100% sequence identity) did not require homology modelling as experimentally determined structures were available. Also, many ADK models were estimated using multiple templates, each of which is shown as a different data point. (B) Distributions of amino acid occurrences at the four regions of Ramachandran plots for all 70 ADKs.

### S3 Relationships between temperature and structural variables

#### S3.1 RMSF

| Model | Phylogenetic random effect | Mean DIC |
|---|---|---|
| RMSF $\sim$ Intercept + $T_{\text{native}}$ + $(T - T_{\text{native}})$ + LID length | ✓ | -544.1021 |
| RMSF $\sim$ Intercept + $(T - T_{\text{native}})$ + LID length | ✓ | -542.0014 |
| RMSF $\sim$ Intercept + $T_{\text{native}}$ + $(T - T_{\text{native}})$ | ✓ | -539.4008 |
| RMSF $\sim$ Intercept + $T_{\text{native}}$ + $(T - T_{\text{native}})$ + ADK type | ✓ | -536.2291 |
| RMSF $\sim$ Intercept + $(T - T_{\text{native}})$ | ✓ | -533.8093 |
| RMSF $\sim$ Intercept + $T_{\text{native}}$ + $(T - T_{\text{native}})$ + LID length | ✗ | -503.0793 |
| RMSF $\sim$ Intercept + $T_{\text{native}}$ + $(T - T_{\text{native}})$ + ADK type | ✗ | -479.0617 |
| RMSF $\sim$ Intercept + $T_{\text{native}}$ + $(T - T_{\text{native}})$ | ✗ | -458.0732 |
| RMSF $\sim$ Intercept + $(T - T_{\text{native}})$ + LID length | ✗ | -449.4954 |
| RMSF $\sim$ Intercept + $T_{\text{native}}$ + LID length | ✓ | -436.7387 |
| RMSF $\sim$ Intercept + $T_{\text{native}}$ + LID length | ✗ | -436.5702 |
| RMSF $\sim$ Intercept + $(T - T_{\text{native}})$ + ADK type | ✗ | -433.3798 |
| RMSF $\sim$ Intercept + $(T - T_{\text{native}})$ | ✗ | -428.3145 |
| RMSF $\sim$ Intercept + $T_{\text{native}}$ + ADK type | ✓ | -427.9029 |
| RMSF $\sim$ Intercept + LID length | ✓ | -427.7656 |
| RMSF $\sim$ Intercept + LID length | ✗ | -426.5244 |
| RMSF $\sim$ Intercept + $T_{\text{native}}$ | ✓ | -423.7988 |
| RMSF $\sim$ Intercept + $T_{\text{native}}$ + ADK type | ✗ | -423.3583 |
| RMSF $\sim$ Intercept + ADK type | ✓ | -418.4271 |
| RMSF $\sim$ Intercept | ✓ | -415.7846 |
| RMSF $\sim$ Intercept + $T_{\text{native}}$ | ✗ | -414.3077 |
| RMSF $\sim$ Intercept + ADK type | ✗ | -414.2151 |
| RMSF $\sim$ Intercept | ✗ | -409.6807 |

Table S2: Candidate models for RMSF, listed from best-fitting to worst-fitting. Models with coefficients whose 95% HPD intervals include 0 are not shown.

##### S3.2 SASA

| Model | Phylogenetic random effect | Mean DIC |
|---|---|---|
| SASA ∼ Intercept + $(T - T_{\text{native}})$ | ✓ | 472.8042 |
| SASA ∼ Intercept + $(T - T_{\text{native}})$ + ADK type | ✓ | 474.1646 |
| SASA ∼ Intercept + $T_{\text{native}}$ + ADK type | ✓ | 480.7414 |
| SASA ∼ Intercept + $(T - T_{\text{native}})$ + LID length | ✓ | 482.2283 |
| SASA ∼ Intercept | ✓ | 486.4846 |
| SASA ∼ Intercept + ADK type | ✓ | 487.4268 |
| SASA ∼ Intercept + LID length | ✓ | 490.9502 |
| SASA ∼ Intercept + $(T - T_{\text{native}})$ + ADK type + LID length | ✗ | 553.6843 |
| SASA ∼ Intercept + $T_{\text{native}}$ + $(T - T_{\text{native}})$ + LID length | ✗ | 557.1076 |
| SASA ∼ Intercept + $(T - T_{\text{native}})$ + LID length | ✗ | 559.2383 |
| SASA ∼ Intercept + $T_{\text{native}}$ + ADK type + LID length | ✗ | 560.4600 |
| SASA ∼ Intercept + $T_{\text{native}}$ + LID length | ✗ | 563.9141 |
| SASA ∼ Intercept + ADK type + LID length | ✗ | 568.7994 |
| SASA ∼ Intercept + LID length | ✗ | 574.5256 |
| SASA ∼ Intercept + $(T - T_{\text{native}})$ + ADK type | ✗ | 711.7906 |
| SASA ∼ Intercept + ADK type | ✗ | 714.7713 |
| SASA ∼ Intercept | ✗ | 724.7923 |

Table S3: Candidate models for SASA, listed from best-fitting to worst-fitting. Models with coefficients whose 95% HPD intervals include 0 are not shown.

##### S3.3 $r_{gyr}$

| Model | Phylogenetic random effect | Mean DIC |
|---|---|---|
| $r_{gyr}$ ∼ Intercept + LID length | ✓ | -372.9937 |
| $r_{gyr}$ ∼ Intercept | ✓ | -370.9737 |
| $r_{gyr}$ ∼ Intercept + $T_{\text{native}}$ + LID length | ✗ | -283.4505 |
| $r_{gyr}$ ∼ Intercept + $(T - T_{\text{native}})$ + LID length | ✗ | -279.3147 |
| $r_{gyr}$ ∼ Intercept + LID length | ✗ | -276.2915 |
| $r_{gyr}$ ∼ Intercept + $T_{\text{native}}$ + ADK type | ✗ | -261.9447 |
| $r_{gyr}$ ∼ Intercept + $(T - T_{\text{native}})$ + ADK type | ✗ | -258.7939 |
| $r_{gyr}$ ∼ Intercept + ADK type | ✗ | -255.8266 |
| $r_{gyr}$ ∼ Intercept | ✗ | -247.2227 |

Table S4: Candidate models for $r_{gyr}$, listed from best-fitting to worst-fitting. Models with coefficients whose 95% HPD intervals include 0 are not shown.

##### S3.4 Number of negatively charged - polar contacts (neg-pol)

| Model | Phylogenetic random effect | Mean DIC |
|---|---|---|
| neg-pol ∼ Intercept + $T_{\text{native}}$ | ✓ | 304.5047 |
| neg-pol ∼ Intercept | ✓ | 305.5843 |
| neg-pol ∼ Intercept + $T_{\text{native}}$ | ✗ | 308.4038 |
| neg-pol ∼ Intercept | ✗ | 316.2293 |

Table S5: Candidate models for the number of negatively charged - polar contacts, listed from best-fitting to worst-fitting. Models with coefficients whose 95% HPD intervals include 0 are not shown.

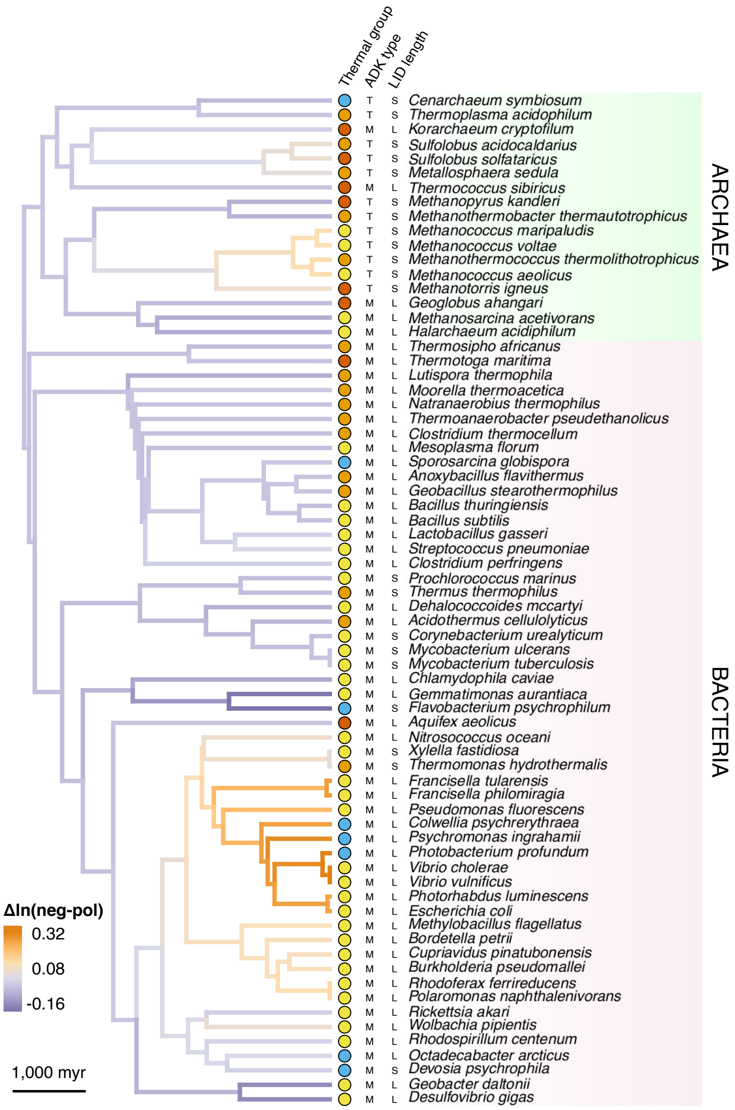

Figure S2: Distribution of the phylogenetic random effect on the intercept of the best-fitting model for negatively charged - polar contact counts (Fig. 8A in the main text) across the branches of the species' phylogeny.

#### S3.5 Number of positively charged - polar contacts (pos-pol)

| Model | Phylogenetic random effect | Mean DIC |
|---|---|---|
| pos-pol ∼ Intercept + ADK type + LID length | ✗ | 214.6132 |
| pos-pol ∼ Intercept + ADK type | ✗ | 216.7225 |
| pos-pol ∼ Intercept + LID length | ✗ | 217.8932 |
| pos-pol ∼ Intercept + ADK type | ✓ | 218.1908 |
| pos-pol ∼ Intercept + LID length | ✓ | 219.4394 |
| pos-pol ∼ Intercept | ✓ | 231.2781 |
| pos-pol ∼ Intercept | ✗ | 237.4459 |

Table S6: Candidate models for the number of positively charged - polar contacts, listed from best-fitting to worst-fitting. Models with coefficients whose 95% HPD intervals include 0 are not shown.

#### S3.6 Number of polar - polar contacts (pol-pol)

| Model | Phylogenetic random effect | Mean DIC |
|---|---|---|
| pol-pol ∼ Intercept + $T_{\text{native}}$ | ✓ | 298.3921 |
| pol-pol ∼ Intercept | ✓ | 303.0924 |
| pol-pol ∼ Intercept + $T_{\text{native}}$ | ✗ | 312.5438 |
| pol-pol ∼ Intercept | ✗ | 313.1715 |

Table S7: Candidate models for the number of polar - polar contacts, listed from best-fitting to worst-fitting. Models with coefficients whose 95% HPD intervals include 0 are not shown.

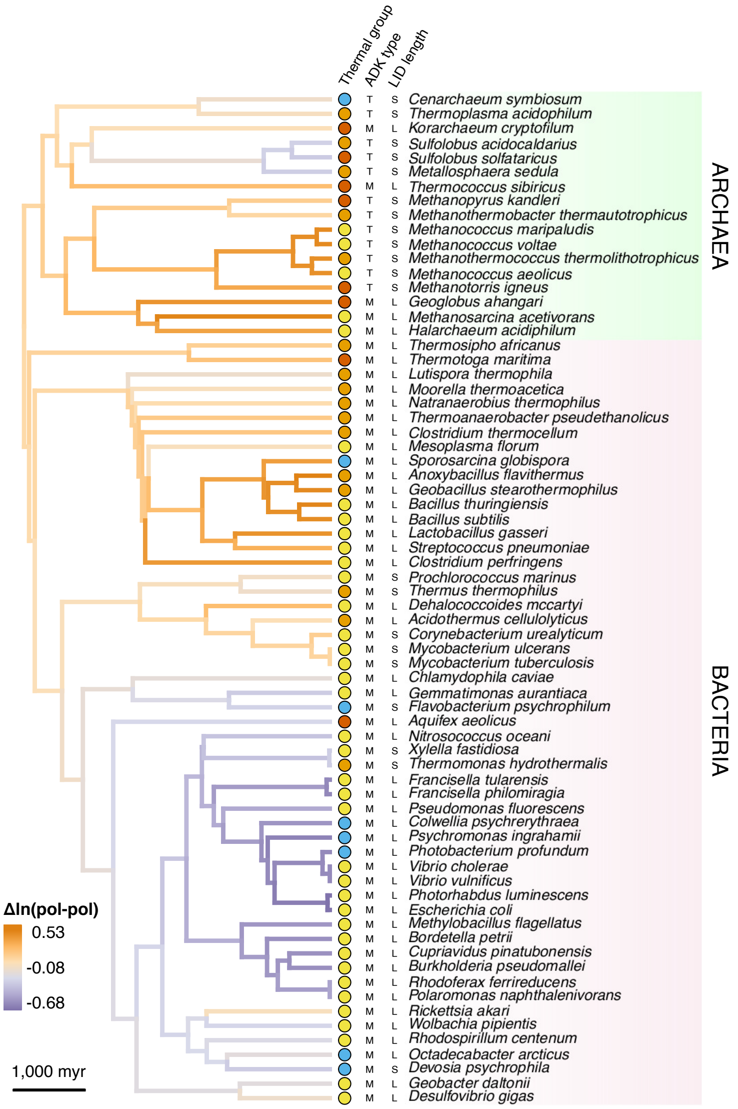

Figure S3: Distribution of the phylogenetic random effect on the intercept of the best-fitting model for polar - polar contact counts (Fig. 8C in the main text) across the branches of the species' phylogeny.

##### S3.7 Number of salt bridges (sb)

| Model | Phylogenetic random effect | Mean DIC |
|---|---|---|
| sb ∼ Intercept + $T_{\text{native}}$ | ✗ | 385.8649 |
| sb ∼ Intercept + $T_{\text{native}}$ | ✓ | 386.5817 |
| sb ∼ Intercept | ✗ | 389.4402 |
| sb ∼ Intercept | ✓ | 389.8820 |

Table S8: Candidate models for the number of salt bridges, listed from best-fitting to worst-fitting. Models with coefficients whose 95% HPD intervals include 0 are not shown.

#### S4 Species-specific relationships between temperature, RMSF, SASA, or $r_{gyr}$

To better compare the effects of temperature on RMSF, SASA, and $r_{gyr}$ among the 5 ADKs for which simulations were performed at 8 temperatures, we fitted variants of the following model with MCMCglmm:

$$\mathrm{Y} = \alpha + \beta \cdot T. \tag{S1}$$

Here, $Y$ is one of RMSF, SASA, or $r_{gyr}$, $\alpha$ is the intercept, and $\beta$ the slope of the relationship. We fitted this model separately to each ADK by executing three Markov chains for 100,000 generations. The first 10,000 generations were treated as burn-in and were removed, following which, posterior samples were obtained every 50 generations. We verified that the three chains had converged to statistically equivalent posterior distributions and that posterior sampling was sufficient, as described in the Methods section of the main text. The resulting estimates are shown in Figs. S4, S5, and S6.

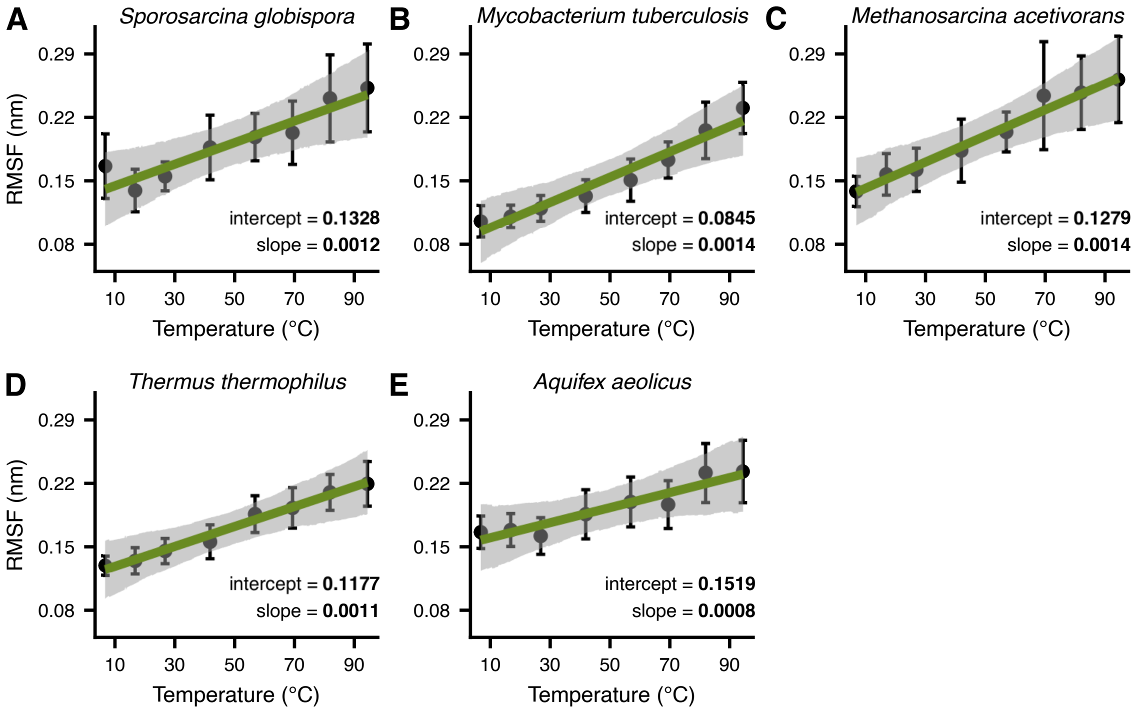

Figure S4: Species-specific relationships between RMSF and temperature. Vertical bars stand for the 95% confidence interval of each data point, whereas grey areas show the 95% Highest Posterior Density (HPD) interval of each fit. Coefficients in bold have a 95% HPD interval that does not include 0.

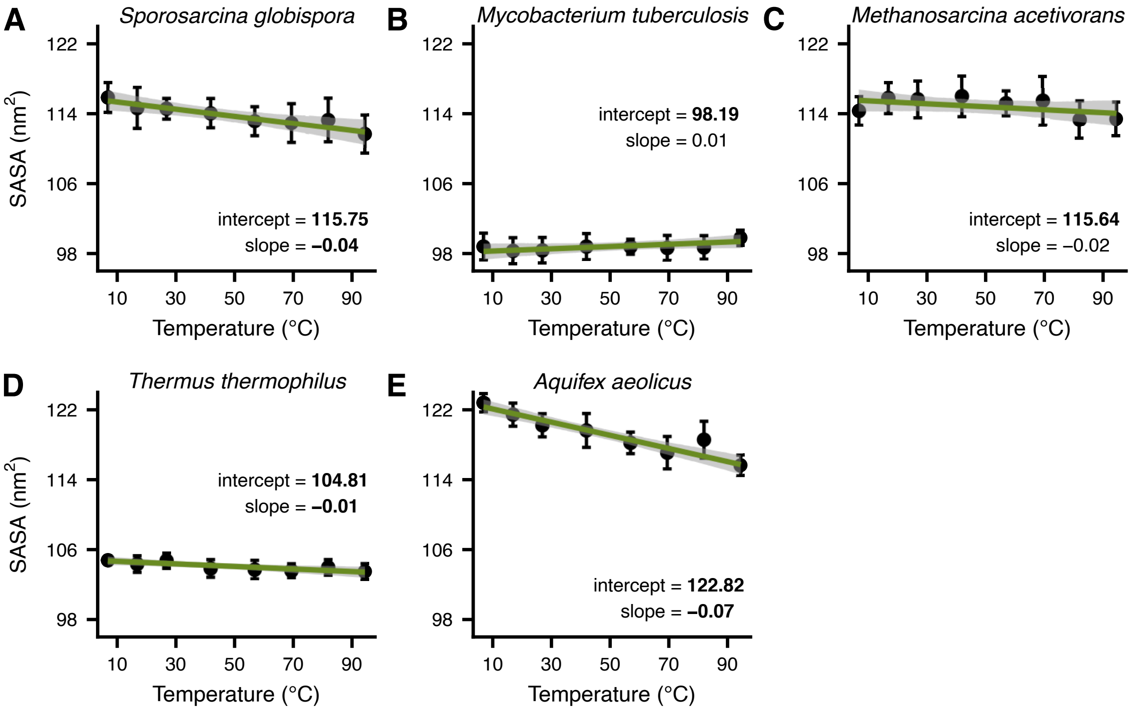

Figure S5: Species-specific relationships between SASA and temperature. Vertical bars stand for the 95% confidence interval of each data point, whereas grey areas show the 95% HPD interval of each fit. Coefficients in bold have a 95% HPD interval that does not include 0.

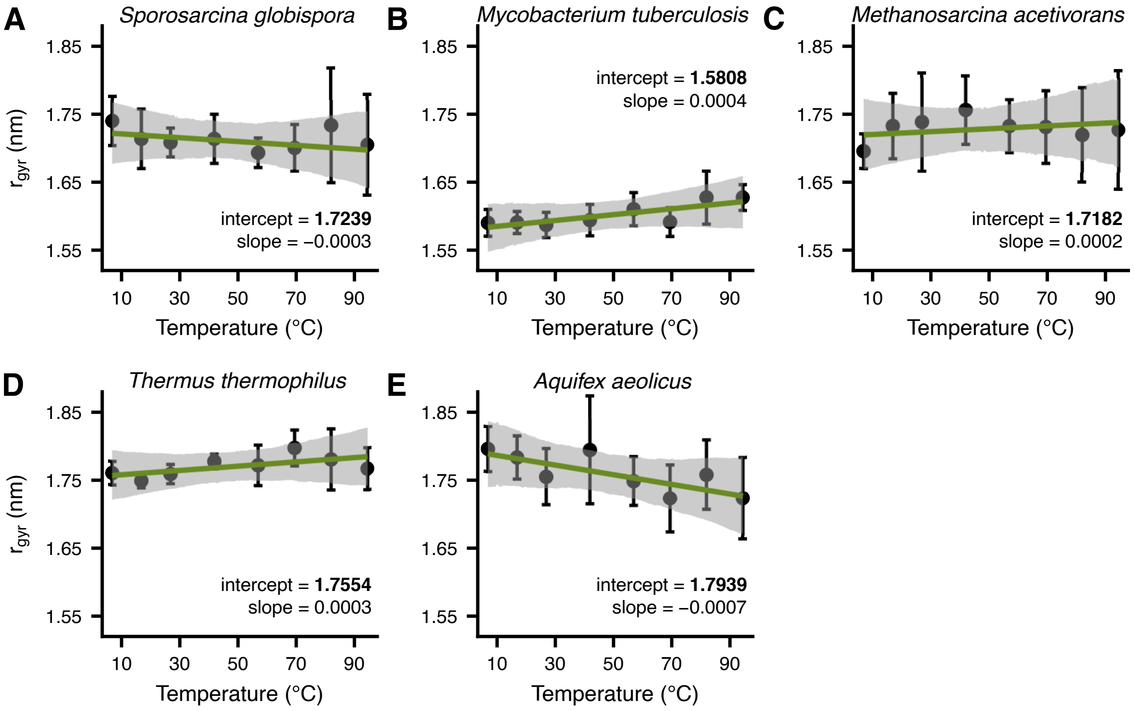

Figure S6: Species-specific relationships between $r_{gyr}$ and temperature. Vertical bars stand for the 95% confidence interval of each data point, whereas grey areas show the 95% HPD interval of each fit. Coefficients in bold have a 95% HPD interval that does not include 0.

#### S5 Average contact networks with a probability of at least 0.7

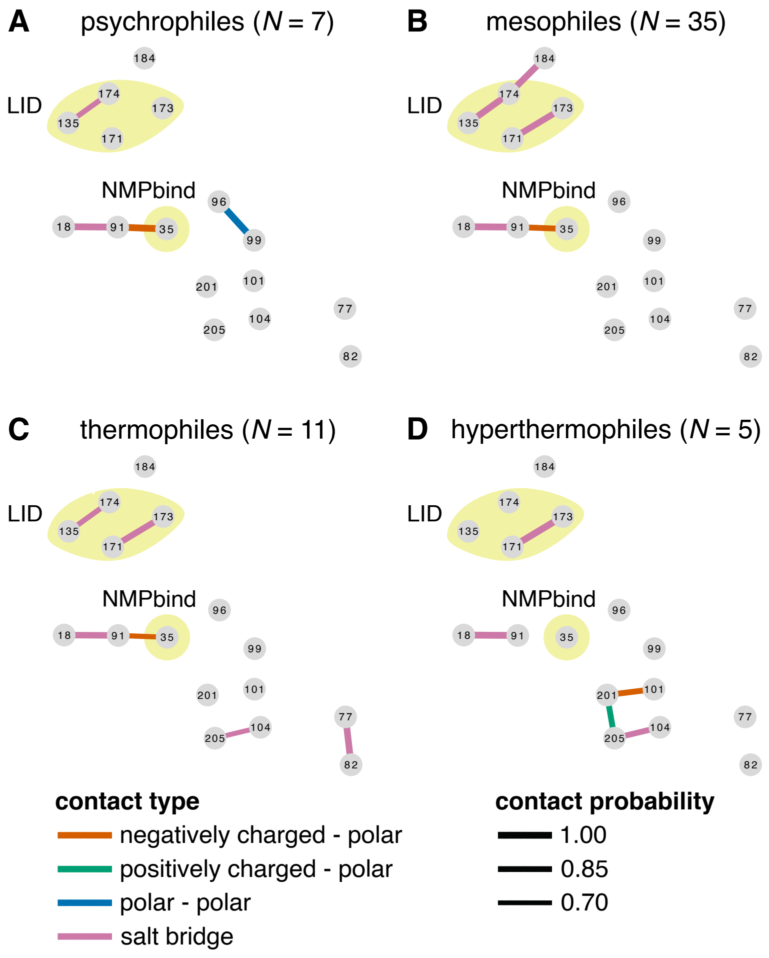

Figure S7: Average contact networks for each thermal group, as in Fig. 9 in the main text. Contacts with a probability below 0.7 are not shown.

#### S6 Correlations between flexibility and kinetic parameters

We examined whether structural flexibility systematically correlates with kinetic parameters ($k_{\text{cat}}$, $K_{\text{M}}$, and $k_{\text{cat}}/K_{\text{M}}$) using the recently released dataset of Muir et al. (2024). Their dataset and ours had 17 species in common. Given that their measurements had been obtained at 25°C, we calculated the RMSF value for each of the 17 species at 25°C based on the model shown in Fig. 5 in the main text, including the species-specific random effect on the intercept. We also extracted the uncertainty around each RMSF estimate from the posterior distribution of our model fit.

To infer the correlations, we fitted three models (one for each kinetic parameter) with MCMCglmm. Each of these models had two covarying response variables (temperature-corrected RMSF and one kinetic parameter), separate intercepts for the two responses, and phylogenetic random effects on the intercepts. Furthermore, the models explicitly accounted for the uncertainty in both the temperature-corrected RMSF and the kinetic parameters. For each model, we executed three independent chains for a million generations, discarding the first 100,000 generations as burn-in, and sampling every 100 generations after that. After performing diagnostics for convergence and sufficient sampling of the posterior (as previously described), we estimated the correlations from the resulting variance-covariance matrices (Fig. S8).

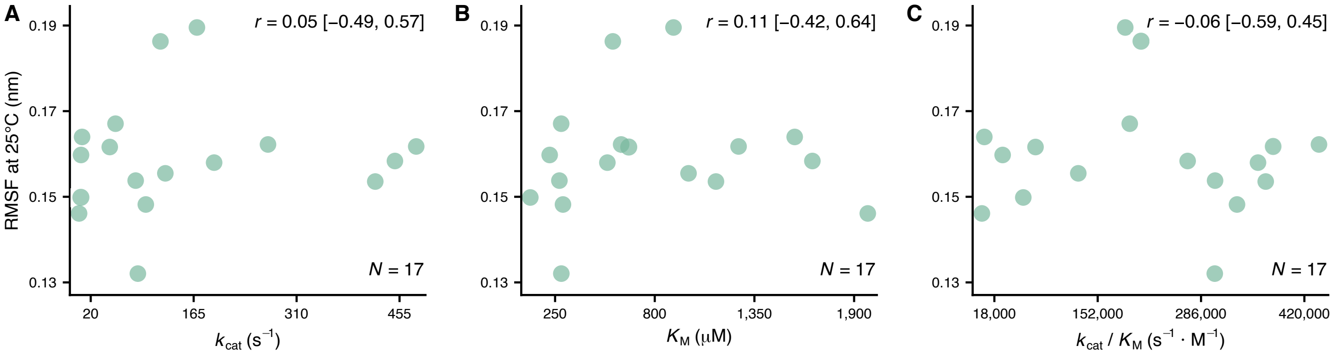

Figure S8: Temperature-normalised structural flexibility appears to be independent of kinetic parameters across 17 prokaryotic ADKs. The reported estimates stand for the mean of the posterior distribution of each correlation and its 95% HPD interval.

### References

Branden, C. I., and J. Tooze. 2012. Introduction to protein structure. Garland Science.

Laskowski, R. A., M. W. MacArthur, D. S. Moss, and J. M. Thornton. 1993. PROCHECK: a program to check the stereochemical quality of protein structures. Journal of Applied Crystallography 26:283–291.

Muir, D. F., G. P. R. Asper, P. Notin, J. A. Posner, D. S. Marks, M. J. Keiser, and M. M. Pinney. 2024. Evolutionary-scale enzymology enables biochemical constant prediction across a multi-peaked catalytic landscape. bioRxiv 2024.10.23.619915.

Rost, B. 1999. Twilight zone of protein sequence alignments. Protein Engineering 12:85–94.
